# Z-RNA-ZBP1 axis drives Tau-mediated neurodegeneration

**DOI:** 10.64898/2026.02.16.706067

**Authors:** Wei Liu, Song-Ang Wu, Bo-Xin Zhang, Shuang-Hui Guo, Lang Li, Wenjing Sun, Xushen Xiong, Jiuhong Nan, Jianfeng Wu, Linghui Zeng, Pilong Li, Zhi-Yu Cai, Huan-Feng Ye, Shuo Zhang, Sheng Nie, Baizhou Li, Dan Wu, Pu Cheng, Xuchen Qi, Dong Fang, Wei Chen, Yingying Zhang, Qiang Chen, Zhang-Hua Yang, Jiahuai Han, Wei Mo

## Abstract

Although the causes of Tau aggregates vary, once Tau aggregates are formed, their neurotoxicity significantly contributes to neuronal death and cognitive decline in tauopathies, with Alzheimer’s disease (AD) being the most well-known example. Despite its central pathogenic role, however, effective therapeutic strategies targeting neurotoxicity of Tau remain poor. Here we demonstrate the pathogenic role of neuronal cell death in Tau-related neurodegeneration. Tau-expressing neurons undergo cell death through Z-DNA-binding protein 1 (ZBP1) activation triggered by endogenous Z-RNAs. These Z-RNAs are derived from reactivated transposable elements (TEs) that are typically silenced within heterochromatin. Tau aggregates show a strong affinity for H3K9me3-modified chromatin, effectively sequestering these epigenetic marks from Heterochromatin Protein 1 (HP1), thereby disrupting the condensation of constitutive heterochromatin. Clinically, an inverse correlation between ZBP1 expression levels in excitatory neurons and cognitive performance in AD patients was observed. Importantly, *Zbp1* haploinsufficiency significantly ameliorated cognitive deficits in aged Tau-transgenic mice (24-month-old), highlighting the therapeutic potential of ZBP1 inhibition to strive against neurodegeneration in tauopathies.

## Main

### Neuronal necroptosis is pathogenic for Tau-mediated neurodegeneration in AD

Tauopathies are a class of neurodegenerative disorders characterized by the abnormal hyperphosphorylation and aggregation of Tau protein, leading to neuronal dysfunction^1^. Representative tauopathies include Alzheimer’s disease (AD)^2^, frontotemporal lobar degeneration^3^ (FTLD), progressive supranuclear palsy (PSP)^4^, and corticobasal degeneration (CBD)^5^. Clinically, these disorders manifest as cognitive impairment, motor dysfunction, or a combination of both, resulting from Tau-induced neuronal dysfunction and loss. Although numerous studies have demonstrated the neurotoxicity of Tau, the precise underlying mechanisms and targeted therapies remain under investigation.

Among tauopathies, Alzheimer’s disease (AD) has attracted the most attention due to its high prevalence and clinical impact. In AD, while amyloid-β (Aβ) is widely considered a key initiator of tau pathology, clinical trials of passive immunotherapy targeting Aβ plaques have failed to preserve cognitive function in patients with established tau pathology^6^. This highlights that once tau aggregates are formed, upstream Aβ clearance alone is insufficient to mitigate the neurotoxic effects of tau. These findings underscore the urgent need to elucidate the mechanisms of Tau-mediated neurotoxicity to enable the development of effective therapeutic strategies targeting Tau pathology directly.

In AD, neuronal death^7^—a defining feature—is increasingly attributed to tau pathology. 5xFAD^8^ and PS19^9^ mouse models were developed to recapitulate Aβ plaque deposition and Tau pathology, respectively. Quantitative analysis of TUNEL staining revealed pronounced neuronal death in the hippocampus and spinal cord of 9-month-old PS19 mice (Extended Data Fig. 1a). In contrast to the pronounced neuronal death observed in tauopathy models, 5xFAD mice showed minimal neuronal loss despite high Aβ burden at comparable ages (Extended Data Fig. 1a). Caspase-8 (Casp8) cleavage and RIPK3 phosphorylation (pRIPK3) serve as critical checkpoints for apoptosis and necroptosis activation, respectively (Extended Data Fig. 1b). To dissect the mechanistic role of Tau aggregate-induced neuronal death in neurodegeneration, we generated PS19 mice with combined *Casp8* and *Ripk3* knockout (PS19, *Casp8*^-/-^, *Ripk3*^-/-^; hereafter PS19, DKO) to block both death pathways (Extended Data Fig. 1c). Intriguingly, although RIPK3 is transcriptionally silenced in healthy neurons, we detected its re-expression in PS19 mouse neurons (Extended Data Fig. 1d, e), suggesting Tau-dependent reactivation of necroptotic machinery.

PS19 mice exhibit progressive cognitive impairment and characteristic motor abnormalities including clasping behavior, limb retraction, and muscle weakness^10^. Genetic blockade of necroptosis through combined deletion of *Casp8* and *Ripk3* in PS19 mice significantly improved spatial cognition (Y-maze test; Fig. 1a) and preserved motor function (grip strength test; Fig. 1b). This intervention also prevented neuronal loss in hippocampal CA1 and CA3 regions (Fig. 1c).

**Fig. 1.**
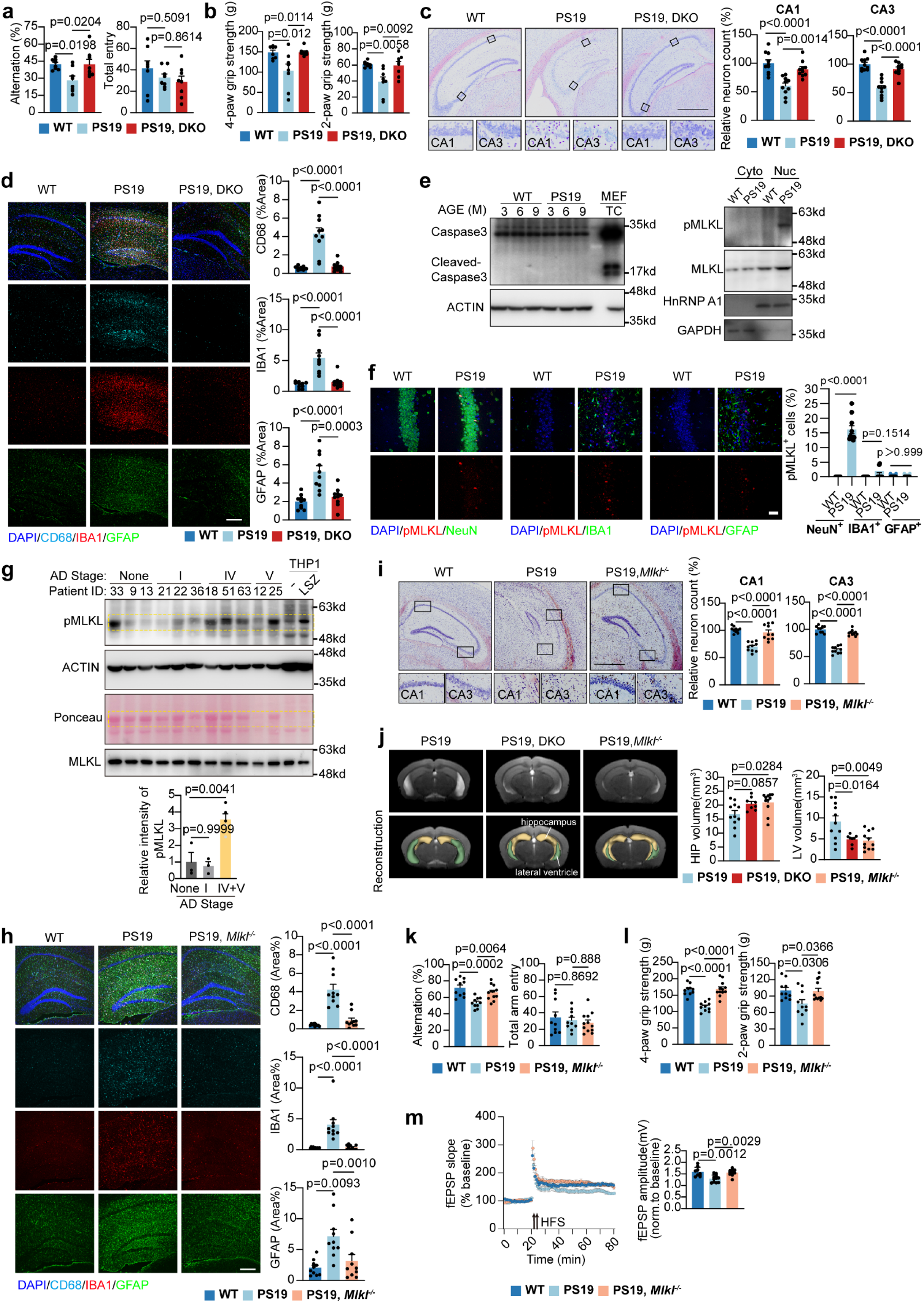
Neuronal necroptosis is pathogenic for Tau-mediated neurodegeneration. **a.** The cognitive function of indicated groups was evaluated by the Y-maze test for measuring total arm entries and spontaneous arm alternation in mice with the indicated genotypes at 9-month-old (n=8 mice). **b.** Quantification of the grip strength in WT, PS19 and PS19, DKO (PS19, *Casp8*^-/-^, *Ripk3*^-/-^) mice at 9-month-old (n=8 mice). **c.** Left: Nissl staining on brain sections of mice with indicated genotypes at 9-month-old. Scale bar: 700 μm. Right: quantification analysis of relative neuron number in CA1 and CA3. n=10 mice. **d.** Representative images and quantification of area of CD68, IBA1 and GFAP signal on brain sections of mice with indicated genotypes. Scale bar: 300 μm. n=10 mice. **e.** Analysis of necroptosis and apoptosis signals in hippocampus of PS19 mice (3-, 6-, 9-month-old) by immunoblotting. Left: Cleaved caspase 3, the marker of apoptosis was not detected in brain samples of PS19 mice. Sample from MEF cells treated with TNF plus cycloheximide (CHX) (TC) was used as positive control for cleaved caspase 3. Right: pMLKL (necroptotic marker) was mainly detected in the nucleus of brain samples of PS19 mice. **f.** Left: representative images of pMLKL co-stained with NeuN, IBA1 or GFAP on brain sections of mice with indicated genotypes at 9-month-old. Scale bar: 20 μm. Right: quantification of percent of pMLKL-positive neurons, IBA1 and GFAP on brain sections. n=10 mice. **g.** Western blots and quantification analysis of the pMLKL signal in hippocampus of AD patients by immunoblotting. Non-AD: n=3, Stage I: n=3, Stage IV+V: n=5. Sample from THP1 cells treated with LPS+SM164+zVAD (LSZ) was used as positive control for pMLKL. **h.** Representative images and quantification of area of CD68, IBA1 or GFAP signal on brain sections of WT, PS19 and PS19, *Mlkl*^-/-^ mice (9-month-old). Scale bar: 300 μm. n= 10 mice. **i.** Left: Nissl staining on brain sections of mice with indicated genotypes at 9-month-old. Scale bar: 700 μm. Right: quantification analysis of relative neuron number in CA1 and CA3. n=10 mice. **j.** Representative MRI images and quantification of brain atrophy (LV volume and HP volume) in mice with indicated genotypes (PS19: 9-month-old, n=11 mice, PS19 DKO, 9-month-old, n=8 mice, and PS19, *Mlkl^-/-^*: 9-month-old, n=5 mice; 11-month-old, n=6 mice). **k.** The cognitive function evaluated by the Y-maze test for measuring total arm entries (right) and spontaneous arm alternation (left) in mice with the indicated genotypes at 9-month-old (WT: n = 10 mice, PS19: n=10 mice, PS19, *Mlkl^-/-^*: n=12 mice). **l.** Quantification analysis of the grip strength in mice with indicated genotypes at 9-month-old (WT, n = 10 mice; PS19, n=10 mice, PS19, *Mlkl^-/-^*: n=12 mice). **m.** LTP recording for continuous 60 min in the hippocampal schaffer collateral region of 9-month-old mice. Averaged potentiation (mean ± s.e.m.) of baseline normalized fEPSP in the indicated groups was calculated (WT: n = 9 slices from 5 mice; PS19: n= 11 slices from 5 mice; PS19, *Mlkl^-/-^*: n=10 slices from 5 mice). Significance between two groups is determined by unpaired *t*-test. Significance between three groups is determined by one way ANOVA test. Data are mean ± s.e.m.

As neuroinflammation represents a hallmark feature of neurodegenerative disease, we examined glial activation in PS19 mice. The double knockout (DKO) mice showed substantial attenuation of microglial and astrocytic activation (Fig. 1d). To characterize the cell death mechanisms, we analyzed hippocampal protein extracts from PS19 mice. While cleaved caspase-3 (apoptosis marker) was undetectable, phosphorylated MLKL (pMLKL, necroptosis marker) was evident at 9 months (Fig. 1e and Extended Data Fig. 1b), indicating predominant necroptotic activity. The pMLKL staining revealed that necroptosis occurred predominantly in neurons rather than glia cells (Fig. 1f). Furthermore, the pMLKL signals localized primarily in neuronal nuclei rather than cytoplasm (Fig. 1f and Extended Data Fig. 1f), consistent with reports showing nuclear MLKL phosphorylation induces milder cell death than classical cytoplasmic necroptosis^11^. This atypical necroptosis phenotype aligns with the chronic neuroinflammation observed in AD. Notably, pMLKL formation remained RIPK3-dependent, as evidenced by its complete absence in PS19, DKO mice (Fig. 1e and Extended Data Fig. 1f). Similarly, phosphorylated RIPK3 detected in 9-month-old PS19 mice (Extended Data Fig. 1g) was abolished in DKO animals. Clinical relevance was confirmed by detecting necroptosis markers in human AD brains, with signal intensity correlating with disease progression (Fig. 1g).

While upregulation of RIPK1 and MLKL is statistically correlated with brain weight loss and cognitive decline in AD patients^12^, the causal role of neuronal necroptosis in neurodegenerative symptoms remains unclear—a question also unresolved in xenograft AD models^13^. Similar to observations in PS19,DKO mice, genetic inhibition of necroptosis via *Mlkl* deletion in PS19 mice significantly attenuated neuroinflammation (Fig. 1h) and neuronal loss (Fig. 1i). Magnetic resonance imaging (MRI) revealed rescue of brain atrophy in both PS19,DKO and PS19,*Mlkl*^-/-^ mice, as evidenced by normalized hippocampal volume and lateral ventricle size (Fig. 1j). Necroptosis blockade further improved spatial cognition (Y-maze test; Fig. 1k), motor function (grip strength test; Fig. 1l) and synaptic plasticity (LTP restoration; Fig. 1m). These findings robustly demonstrate that necroptosis-mediated neuronal death drives tau-induced neurodegeneration, prompting mechanistic investigation of this pathway.

### Tau aggregates induce ZBP1-dependent neuronal necroptosis

To determine whether neuronal necroptosis results from pathogenic tau aggregate accumulation, we established an *in vivo* Tau acceleration model. Briefly, hippocampal injection of tau aggregates in 2.5-month-old PS19 mice significantly accelerated tau propagation within 3 weeks (hereafter referred to as the Tau acceleration model; Extended Data Fig. 2a). Pathological tau was identified using AT8 (pS202/pT205) immunostaining (Extended Data Fig. 2b, red). We further confirmed phosphorylation at additional disease-relevant sites including T181, T231^14^, S356^15^, and S396^16^ (Extended Data Fig. 2b), demonstrating that Tau acceleration injection induced concurrent hyperphosphorylation at multiple epitopes (Extended Data Fig. 2b).

In 9-month-old PS19 mice, AT8-positive tau inclusions were consistently co-labeled with antibodies against pT181, pT231 and pS396 (Extended Data Fig. 2c), while approximately 20% of AT8-positive aggregates showed pS356 immunoreactivity (Extended Data Fig. 2c). X-34 staining confirmed the β-sheet conformation characteristic of mature aggregates (Extended Data Fig. 2d). The accumulation of Tau aggregate in Tau acceleration model penocopied PS19 mice (Extended Data Fig. 2d, e), and these Tau aggregates also induced pronounced necroptosis (pMLKL) (Extended Data Fig. 2f) and neuroinflammation similar to that seen in 9-month-old PS19 mice (Extended Data Fig. 2g).

To further confirm that pathological Tau directly induces neuronal death, *in vitro* culture system was used. We cultured primary cortical neurons from the cortex of newborn PS19 mice (Extended Data Fig. 3a). Previous studies have demonstrated that extracellular tau aggregates are internalized into neurons via the low-density lipoprotein receptor-related protein 1 (LRP1)^17^. Primary cortical neurons incubated with recombinant human Tau aggregates but not WT Tau protein exhibited immunopositivity for both phosphorylated tau (AT8 epitope) and β-sheet-rich conformations, as detected by the amyloid dye X-34 (Fig. 2a and Extended Data Fig. 3b).

**Fig. 2.**
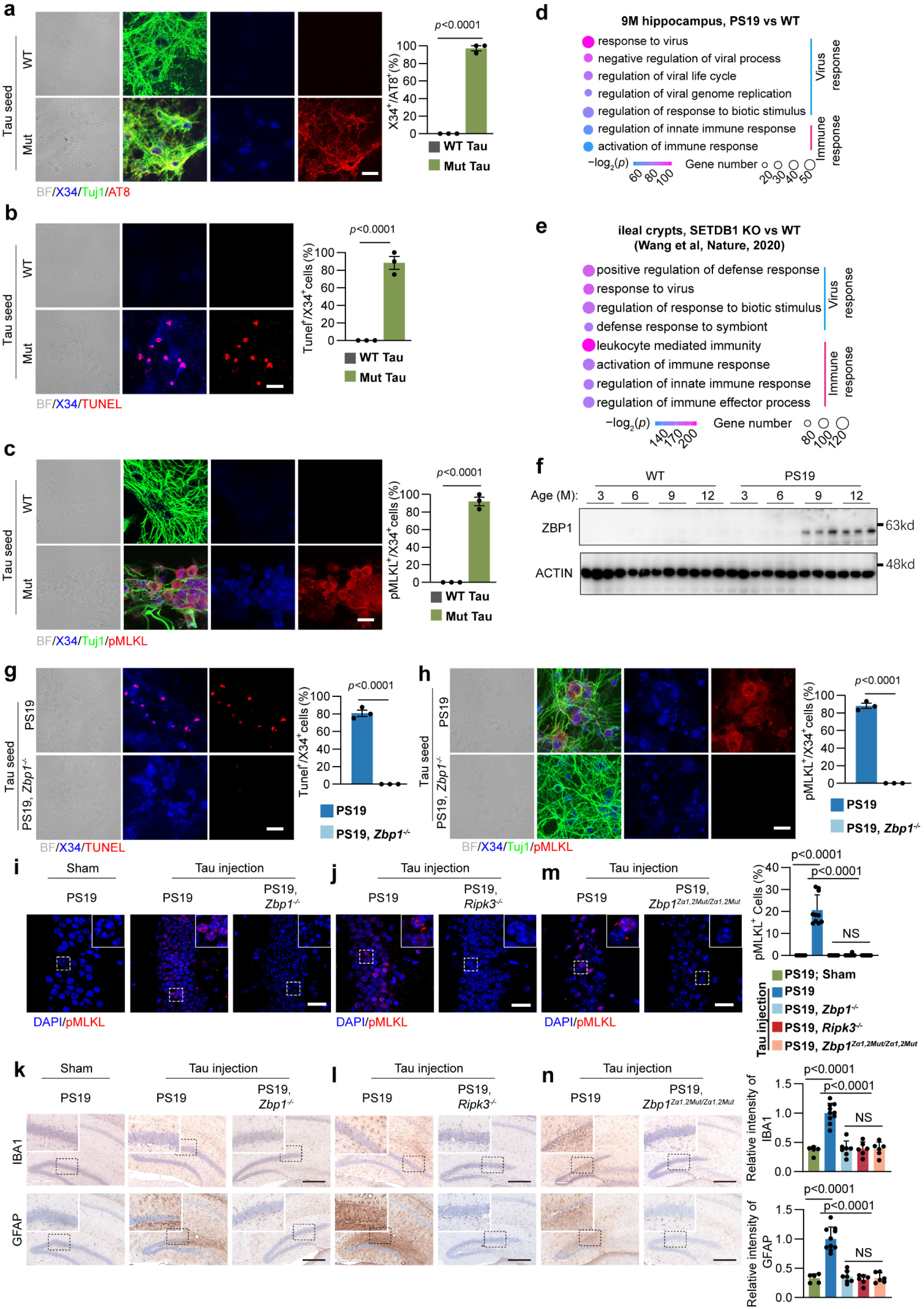
Tau aggregate generates death ligand to active ZBP1 for necroptosis. **a.** Representative images of X34, AT8 and Tuj1 staining and quantification of AT8^+^/X34^+^ cells for cortical neurons from PS19 mice treated with recombinant WT or mutant Tau protein. Scale bar: 20 μm. n=3 independent experiments. **b.** Representative images of Tunel and X34 staining and quantification of Tunel^+^/X34^+^ cells for cortical neurons from PS19 mice treated with recombinant WT or mutant Tau protein. Scale bar: 20 μm. n=3 independent experiments. **c.** Representative images of X34, pMLKL and Tuj1 staining and quantification of pMLKL^+^/X34^+^ cells for cortical neurons from PS19 mice treated with recombinant WT or mutant Tau protein. Scale bar: 20 μm. n=3 independent experiments. **d.** Global transcriptome analysis of hippocampus from 9-month-old mice (PS19 versus WT). **e.** Global transcriptome analysis of ileal crypts (SETDB1 KO versus WT) from published work (Wang et al, Nature, 2020). **f.** Immunoblotting of ZBP1 protein level in hippocampus of WT and PS19 mice at indicated age (n=3 mice). M: month. **g.** Representative images of Tunel and X34 staining and quantification of Tunel^+^/X34^+^ cells for cortical neurons from PS19 and PS19, *Zbp1*^-/-^ mice treated with recombinant mutant Tau protein. Scale bar: 20 μm. n=3 independent experiments. **h.** Representative images of X34, Tuj1 and pMLKL staining and bright filed and quantification of pMLKL^+^/X34^+^ cells for cortical neurons from PS19 and PS19, *Zbp1*^-/-^ mice treated with recombinant mutant Tau protein. Scale bar: 20 μm. n=3 independent experiments. **i,j,m.** Representative images and quantification analysis of percent of neuronal death (pMLKL-positive) in Tau acceleration model with indicated genotypes (Sham PS19: n=5 mice, PS19: n= 10 mice; PS19, *Zbp1^-/-^*: n=7 mice; PS19, *Ripk3^-/-^*: n=6 mice; PS19, *Zbp1^Za^*^1,^*^2Mut/Za^*^1,^*^2Mut^*: n=6 mice). Selected areas are shown magnified to the right of each image. Scale bar: 20 μm. **k.l.n.** Representative images and quantification analysis of neuroinflammation (relative intensity of IBA1 or GFAP signal) in Tau acceleration model with indicated genotypes (Sham PS19: n=7 mice, PS19: n= 10 mice; PS19, *Zbp1^-/-^*: n=7 mice; PS19, *Ripk3^-/-^*: n=6 mice; PS19, *Zbp1^Za^*^1,^*^2Mut/Za^*^1,^*^2Mut^*: n=6 mice). Selected areas are shown magnified to the right of each image. Scale bar: 300 μm. Significance between two groups is determined by unpaired *t*-test. Significance between three groups is determined by one way ANOVA test. Data are mean ± s.e.m. NS, not significant.

Furthermore, these neurons showed elevated levels of TUNEL (Fig. 2b) and phosphorylated MLKL (pMLKL) (Fig. 2c), suggesting that direct uptake of Tau aggregates induces neuronal necroptotic cell death.

In other tauopathies, Tau pathology is also known to induce neuronal death and loss. To investigate whether tau aggregates can trigger cell death in other neuronal subtypes, we utilized cultured dorsal root ganglion (DRG) neurons from PS19 mice to further elucidate the pathological mechanisms of tauopathy^18^ (Extended Data Fig. 3c). We exposed DRG neurons isolated from 2-month-old PS19 mice to the same Tau aggregates (Extended Data Fig. 3c, d). Consistent with our findings in cortical neurons, internalized tau aggregates directly induced neuronal necroptosis in DRG neurons (Extended Data Fig. 3c, d).

While classical necroptosis is typically mediated by extracellular death ligands such as TNF (Extended Data Fig. 3e, left half), we sought to determine whether autocrine TNF contributed to tau-induced neuronal death. TNF neutralization experiments using anti-TNF antibodies demonstrated that Tau aggregate-induced neuronal death was not attenuated by TNF blockade (Extended Data Fig. 3f), indicating a TNF-independent mechanism.

To elucidate the molecular pathways underlying TNF-independent necroptosis, we conducted RNA sequencing to compare hippocampal transcriptomes between 9-month-old PS19 and wild-type (WT) mice. The most significantly upregulated genes in PS19 hippocampus were associated with viral infection and antiviral responses (Fig. 2d and Extended Data Fig. 3g). Notably, this transcriptional profile closely resembled that of intestinal epithelial cells undergoing ZBP1-dependent necroptosis, as we previously reported (Extended Data Fig. 3e, right half; Fig. 2e and Extended Data Fig. 3h).

ZBP1 (Z-DNA binding protein 1) is an interferon-stimulated gene (ISG)^19^. In healthy brains, ZBP1 protein expression is barely detectable. However, in PS19 mice, ZBP1 was robustly upregulated by 9 months of age (Fig. 2f), coinciding with the onset of necroptosis. Of note, the ZBP1 staining in PS19 mice revealed its predominant expression in neurons rather than glia cells (Extended Data Fig. 3i). In humans, ZBP1 exists as two isoforms: a 429-amino acid (aa) variant and a 354-aa variant (Extended Data Fig. 3j). Under basal conditions, ZBP1 is present in normal aging brains, likely due to age-related chronic inflammation. Notably, ZBP1 expression was moderately elevated in early-stage Alzheimer’s disease (AD) patients and further increased in advanced stages (Extended Data Fig. 3k). These clues prompt us to speculate whether Tau-aggregate-induced neuronal death is mediated by ZBP1.

Importantly, in primary PS19 neurons, including both cortical neurons and DRG neurons, *Zbp1* deletion effectively blocked Tau aggregate-induced necroptosis *in vitro* (Fig. 2g, 2h and Extended Data Fig. 3l). Similarly, in the *in vivo* Tau acceleration model, Tau aggregate-induced necroptosis in hippocampal neurons was abolished by either *Zbp1* knockout or deletion of its downstream effector *Ripk3* (Fig. 2i, 2j and Extended Data Fig. 3m, 3n). These findings establish that Tau aggregates trigger ZBP1-dependent necroptotic neuronal death. Notably, Tau aggregate-induced neuroinflammation was also significantly attenuated in the hippocampus of *Zbp1*- or *Ripk3*-deficient PS19 mice (Fig. 2k, 2l).

Under most conditions, ZBP1-mediated necroptosis requires activation by dsRNA or dsDNA (Extended Data Fig. 3e, right panel). This activation occurs through binding of ZBP1’s Zα domains (Zα1 and Zα2) to these nucleic acids^20,21^. To determine whether dsRNA/dsDNA is necessary for ZBP1-dependent cell death in tauopathy, we generated PS19 mice with ZBP1 Zα1/2 mutations^22^ (containing four amino acid substitutions that abrogate nucleic acid binding). Remarkably, Tau-induced neuronal death and neuroinflammation were completely rescued in PS19, ZBP1 Zα1/2 mutant mice (Fig. 2m, 2n and Extended Data Fig. 3o), demonstrating that Tau aggregates cannot directly initiate ZBP1-dependent necroptosis, but rather require endogenous production of dsRNA or dsDNA as obligatory death ligands.

To facilitate the investigation of the mechanisms underlying Tau aggregate-induced neuronal cell death, we established an *in vitro* cellular tauopathy model using cell lines rather than primary cultured neurons, which provides a tractable and reproducible system to mimic Tau aggregate accumulation in neurons. This model was originally established in 293T cells^23^. We replaced 293T cells with neuroblastoma SH-SY5Y cells, as they can differentiate into neurons and are widely used to study the molecular mechanisms underlying neurodegenerative diseases^24,25^. Cells were expressed with Tau-4R (242-372aa of the 441aa Tau 2N4R, P301L mutated; named Tau-4R cells) (Extended Data Fig. 4a), the aggregation-competent core of Tau^26^. Tau puncta was observed in Tau-4R cells transfected with Tau seeds derived from brain protein lysates of rTg4510 mice (Extended Data Fig. 4a). All Tau puncta was phosphorylated at Ser 356 site (Extended Data Fig. 4b), a site previously identified as a pathological phosphorylation marker in Tau accelerated model (Extended Data Fig. 2b). Furthermore, Tau puncta was shown positive for X34^27^ (Extended Data Fig. 4c) and Thio-S^28^ (Extended Data Fig. 4d), two well-established markers of Tau aggregates, confirming that these puncta represent accumulated Tau aggregates. Notably, ∼ 70% of Tau-4R cells containing Tau puncta underwent necroptosis, whereas puncta-negative Tau-4R cells showed no signs of necroptosis (Extended Data Fig. 4e), directly linking Tau aggregates to necroptotic cell death. In summary, this cellular system provides a robust model for studying the mechanisms by which Tau aggregation drives neuronal death.

### Z-RNAs from reactivated TEs bind to ZBP1

Next, we sought to identify endogenous nuclei acids that activate ZBP1 in PS19 mice. We previously reported that ZBP1 sensed self-dsRNA to activate necroptosis in *Setdb1*-deleted gut^19^. *Setdb1*-deleted organiod^19^ as well as cells transfected with poly (I:C) were used as the positive control of dsRNA staining visualized by J2 antibody (Extended Data Fig. 5a, 5b). In Tau-4R cells, about 90% Tau puncta cells were positive for J2 staining, which was resistant to DNase digestion but vanished by RNase digestion (Extended Data Fig. 5c). In 9-month-old PS19 mice, J2 signal was detected in hippocampal granule neurons (Extended Data Fig. 5d).

Under virus-free conditions, the dsRNAs accumulated in neurons are of endogenous origin. As demonstrated in our previous study^19^, these endogenous dsRNAs primarily derive from reactivated endogenous retrovirus (ERV) transcripts, which belong to the long terminal repeat (LTR) class of transposable elements (TEs). To determine the source of dsRNAs in PS19 mice, we analyzed hippocampal long non-coding RNA sequencing (lncRNA-seq) data. This analysis confirmed robust TE activation (Fig. 3a) and revealed a significant upregulation of several TE families, notably LTR and LINE elements, in 9-month-old PS19 hippocampal neurons (Fig. 3b).

**Fig. 3.**
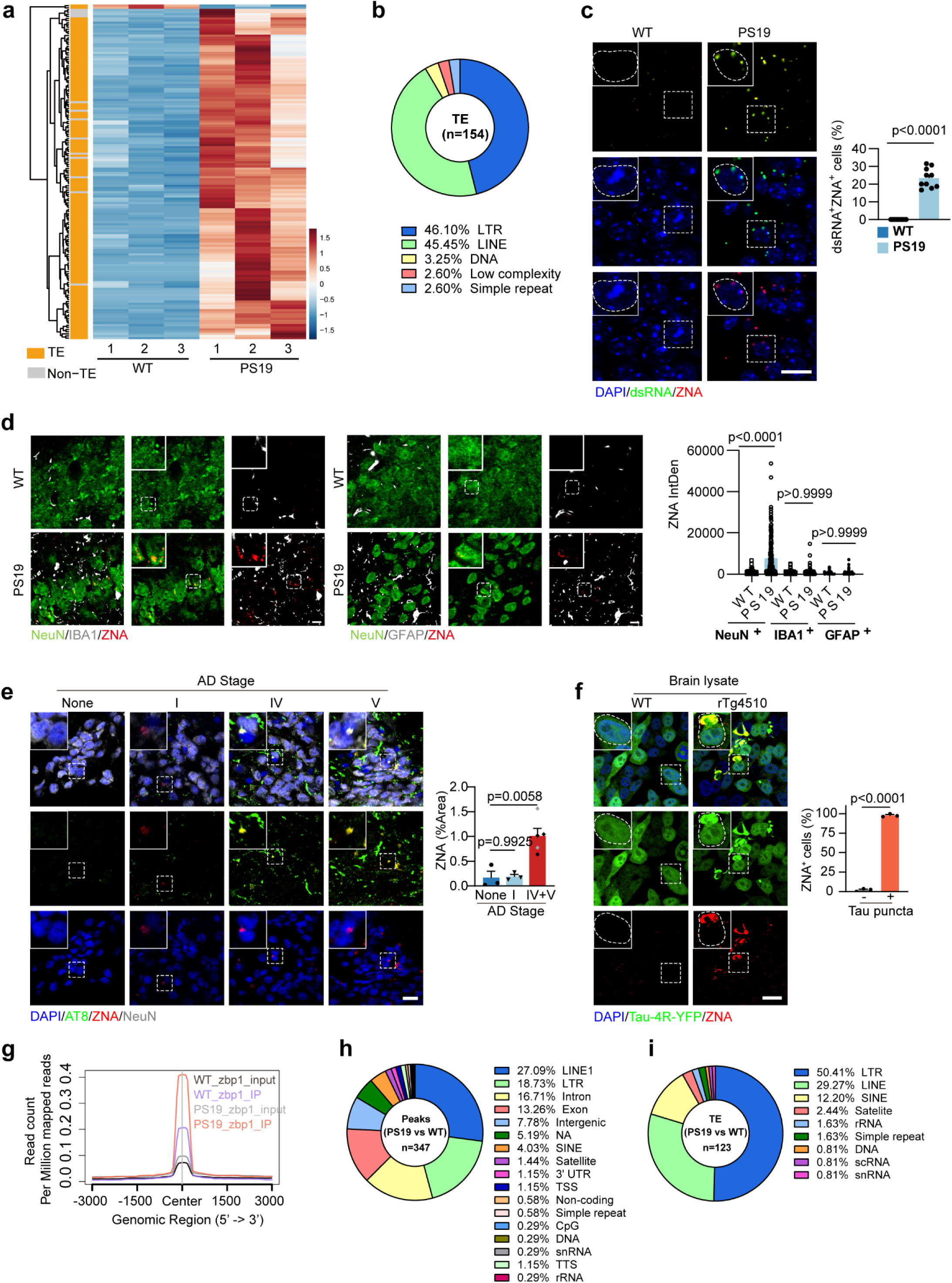
Pathogenic tau produces Z-RNA from reactivated TEs. **a.** Heatmap of upregulated non-coding RNA (PS19 vs WT, fold change>4, P<0.05) in hippocampus of WT and PS19 mice at 9-month-old (n = 3) by LncRNA-seq. **b.** Classification of TEs unregulated in PS19. **c.** Left: representative images of dsRNA (green) and ZNA (red) staining on brain sections of WT and PS19 mice at 9-month-old. Scale bar: 10 μm. Selected areas are shown magnified to the left of each image. Right: quantification of percent of dsRNA^+^ZNA^+^ cells. **d.** Left: representative images of ZNA (red) co-stained with NeuN, IBA1 or GFAP on brain sections of mice with indicated genotypes at 9-month-old. Scale bar: 10 μm. Right: quantification of intensity of ZNA in neurons, IBA1 and GFAP with indicated genotypes (n=10 mice). **e.** Left: representative images of ZNA (red) and phosphorylated tau (AT8, green) staining in hippocampus of AD patients at various stages. Scale bar: 20 μm. Selected areas are shown magnified to the left of each image. Right: quantification analysis of the ZNA signal (%Area) in hippocampus of AD patients. Non-AD: n=3, Stage I: n=3, Stage IV+V: n=5. **f.** Left: representative images of ZNA (red) staining on SH-SY5Y (Tau-4R-YFP) cells treated with brain lysates from rTg4510 or WT mice. Scale bar: 20 μm. Selected areas are shown magnified to the left of each image. Right: quantification of the percent of ZNA^+^ cells (%) in Tau puncta^+^ cells. n=3 independent experiments. **g.** Metaplot of RNA immunoprecipitation (RIP)-RNA signals in the significantly up-regulated peak center (log2FC (IP/input) > 0, *p* < 0.05) (n = 347). **h.** Pie charts showing the distributions of upregulated peaks identified by RNA immunoprecipitation sequencing (RIP-seq) of ZBP1 (log2FC (PS19/WT) > 0, *p* < 0.05) (n = 347). **i.** Classification of TEs significantly unregulated in PS19 by ZBP1-RIP. Significance between two groups is determined by unpaired *t*-test. Significance between three or more groups is determined by one way ANOVA test. Data are mean ± s.e.m.

dsRNA recognized by ZBP1 exhibits distinct structural features compared to RNA sensed by other RNA sensors. Unlike right-handed B-DNA and A-RNA, Z-form nucleic acids (ZNA, including Z-RNA and Z-DNA) adopt a left-handed conformation characterized by a zigzag backbone^29,30^. ZNA is thermodynamically unstable both *in vitro* and *in vivo*^29^.

In PS19 mice, a significant proportion of neuronal dsRNA adopted the Z-form, as visualized by ZNA-specific antibody staining (Fig. 3c). Consistent with the observation that necroptosis occurs predominantly in neurons (Fig. 1f), ZNA signals were also enriched in neurons rather than glial cells (Fig. 3d). The presence of Z-RNA in neurons was further validated by DNase/RNase treatment (Extended Data Fig. 5e). Moreover, Z-RNA was detected in postmortem brain tissues from Alzheimer’s disease (AD) patients (Fig. 3e) and confirmed by DNase/RNase sensitivity assays (Extended Data Fig. 5f).

Furthermore, we demonstrated that Z-RNA formation is directly linked to Tau aggregation. In Tau-4R cells, ∼90% of Tau puncta-positive cells exhibited Z-RNA accumulation, whereas no Z-RNA signal was detected in Tau puncta-negative cells (Fig. 3f), indicating a strict correlation between Tau aggregate formation and Z-RNA generation.

To identify the Z-RNAs that bound to ZBP1, RNA immunoprecipitation and sequencing (RIP-seq) were performed using a ZBP1-specific antibody (Extended Data Fig. 5h). For more precise characterization of ZBP1-recognized Z-RNAs, we used WT mouse neurons as a control to determine sequences specifically enriched by ZBP1 in PS19 mouse neurons (Fig. 3g and Extended Data Fig. 5h). By comparing ZBP1-immunoprecipitated RNAs to total input RNA, we found that nearly half (49.85%) of the RNAs enriched in PS19 neurons were TEs, primarily composed of LINE1 (27.09%), LTR (18.73%) and SINE elements (4.03%) (Fig. 3h, 3i).

### Pathogenic Tau deforms heterochromatin condensation

TEs are primarily silenced in constitutive heterochromatin, which is marked by H3K9me3^31–34^. In PS19 mice, the expression of SETDB1 (a histone H3K9 methyltransferase responsible for H3K9me3 deposition) remained unchanged (Extended Data Fig. 6a). Constitutive heterochromatin is typically localized near the nuclear lamina^31,33–36^.

In hippocampal neurons of PS19 mice, nuclear lamina wrinkling or invagination was not observed until 9 months of age, coinciding with the accumulation of pathological Tau (Fig. 4a). The proportion of neurons exhibiting abnormal nuclear lamina morphology increased further at 10 months (Fig. 4a). Consistent with these structural changes, H3K9me3 levels were significantly reduced starting at 9 months (Fig. 4b), suggesting progressive heterochromatin relaxation in Tau-affected neurons.

**Fig. 4.**
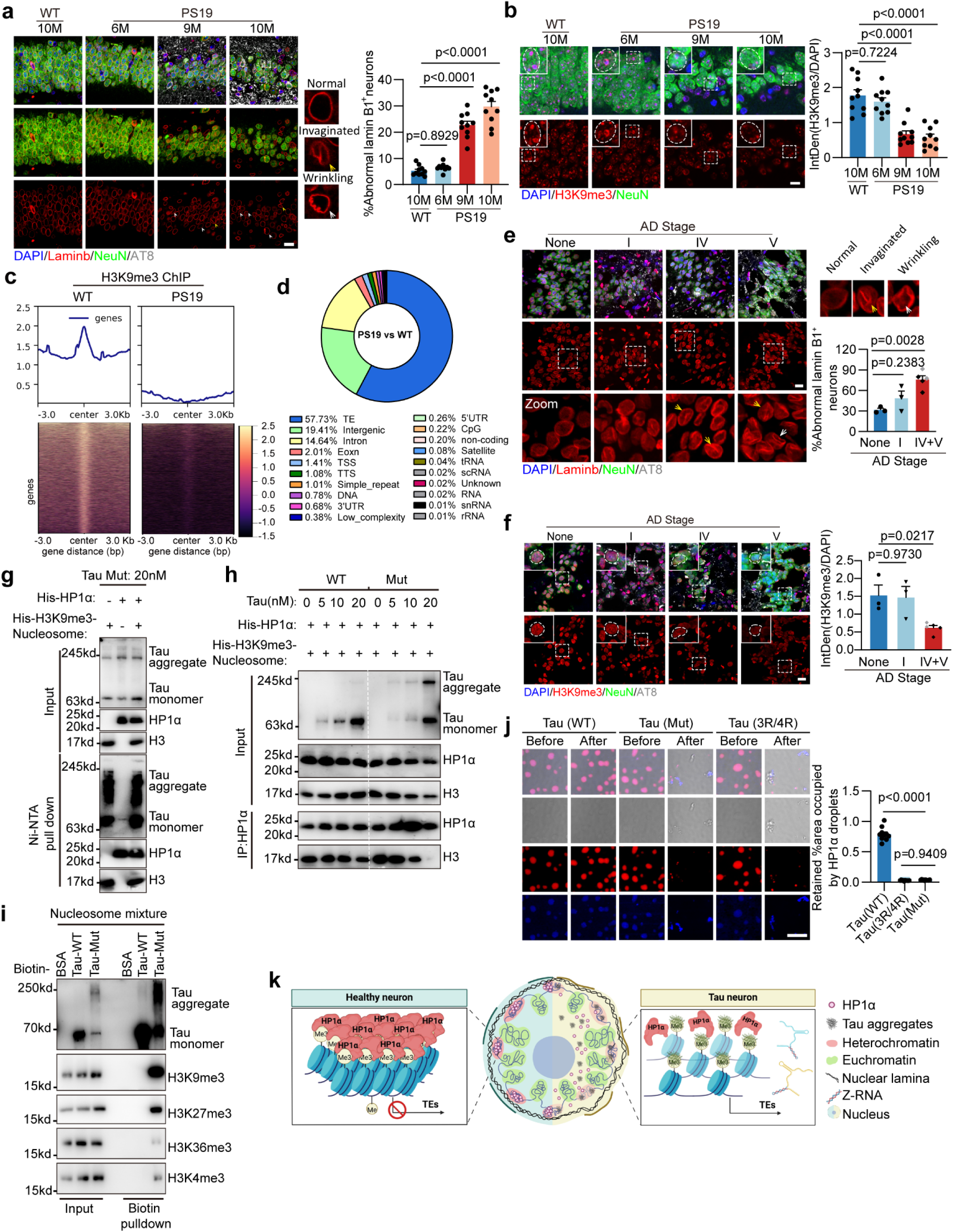
Tau aggregate breaks the condensation of constitutive heterochromatin. **a.** Left: representative images of Lamin B1, AT8 and NeuN staining in hippocampus of WT and PS19 mice at 6-, 9- and 10-month-old. M: month. Right: quantification analysis of percent of neurons with abnormal Lamin B1. Scale bar: 10 μm. n=10 mice. **b.** Left: representative images of H3K9me3 and NeuN staining in hippocampus of WT and PS19 mice at stages indicated. Right: quantification analysis of relative intensity of H3K9me3/DAPI. Scale bar: 10 μm. n=10 mice. **c.** H3K9me3-binding sites shown in heatmap. ±3 kb surrounding H3K9me3-binding summits. **d.** The distribution of peaks with 2-fold downregulation of *p*>0.05 in H3K9me3 ChIP in hippocampus of PS19 compared with WT control. **e.** Left: representative images of Lamin B1, p-Tau (AT8) and NeuN staining on hippocampus sections of AD patients at various stages as indicated. Scale bar: 10 μm. The representative images for normal, invaginated and wrinkling nuclear membrane were shown in the right upper. Right lower: quantification of abnormal Lamin B1^+^ neurons (%). Non-AD: n=3, Stage I: n=3, Stage IV+V: n=5. **f.** Left: representative images of H3K9me3, p-Tau and NeuN staining on hippocampus sections of AD patients at various stage as indicated. Scale bar: 10 μm. Right: quantification of relative intensity of H3K9me3/DAPI. Non-AD: n=3, Stage I: n=3, Stage IV+V: n=5. **g.** Recombinant mutant tau protein (20 nM) was incubated with His-tagged HP1α or H3K9me3-marked nucleosome respectively. Inputs and Ni-NTA pull-down samples were analyzed by immunoblotting as indicated. H3: H3K9me3-marked histone 3. **h.** Various doses of WT or mutant Tau protein was added to the pre-incubated His-tagged HP1α and H3K9me3-marked nucleosome. The inputs and anti-HP1α immunoprecipitates were analyzed by immunoblotting as indicated. **i.** Recombinant mutant tau protein labeled with biotin was incubated with nucleosome mixture including equivalent H3K9me3-nucleosome, H3K27me3-nucleosome, H3K36me3-nucleosome and H3K4me3-nucleosome. Inputs and Biotin pull-down samples were analyzed by immunoblotting as indicated. **j.** Left: representative images of droplets formation by LLPS of HP1α with H3K9me3-marked nucleosomal arrays (NA) treated with recombinant WT, mutant tau protein and Tau (3R/4R) (3 μM). Scale bar: 10 μm. Right: quantification of retained area (%) occupied by HP1α droplets. n=10 independent experiments. **k.** Proposed model for disruption of heterochromatin condensation by pathogenic tau to generate death ligand to active ZBP1-dependent necroptosis. Significance between three or more groups is determined by one way ANOVA test. Data are mean ± s.e.m.

Critically, ChIP-seq analysis revealed a genome-wide reduction in H3K9me3 enrichment at multiple TE loci in PS19 hippocampal neurons (Fig. 4c, 4d), providing direct evidence for TE reactivation.

Xiong et al. established a single-cell epigenomic atlas of the prefrontal cortex (PFC) and demonstrated progressive epigenetic erosion in late-stage Alzheimer’s disease (AD) patients^37^. Strikingly, our findings in mouse models were strongly corroborated in human AD pathology. We observed severe constitutive heterochromatin erosion in hippocampal neurons of AD patients, which showed a close spatial association with neurofibrillary tangles (NFTs). Notably, in regions containing Tau aggregates, we detected distorted nuclear architecture, including wrinkled or invaginated nuclear lamina (Fig. 4e). Consistent with the mouse data, we found that H3K9me3 modification levels were significantly reduced in late-stage AD patients (Fig. 4f), further supporting the conserved role of heterochromatin destabilization in Tau-driven neurodegeneration.

To determine whether Tau aggregate accumulation disrupts heterochromatin organization, we employed our *in vitro* cellular tauopathy model. Strikingly, Tau aggregate induction caused significant nuclear lamina deformation. While 85% of Tau puncta-positive cells exhibited abnormal nuclear lamina morphology, only 19% of puncta-negative cells showed similar defects (Extended Data Fig. 6b).

In Tau puncta-positive cells, we observed moderate reductions in both the repressive histone mark H3K9me3 and its epigenetic reader protein HP1α (Extended Data Fig. 6c). Given that HP1α binding to H3K9me3-modified nucleosomes is essential for heterochromatin maintenance^38^, the significantly decreased co-localization frequency between HP1α and H3K9me3 in puncta-positive cells (Extended Data Fig. 6c) strongly suggests that Tau aggregates specifically disrupt HP1α recruitment to H3K9me3-marked chromatin regions.

To investigate whether and how Tau aggregates impair HP1α recruitment to H3K9me3-modified chromatin, we generated recombinant HP1α proteins and reconstituted H3K9me3-nucleosomes for cell-free biochemical studies (Extended Data Fig. 6d). Pulldown assays demonstrated that 20 nM Tau aggregates (but not wild-type Tau) abolished HP1α binding to H3K9me3-nucleosomes (Extended Data Fig. 6e), indicating direct interference with this epigenetic interaction. To further elucidate the mechanism of this disruption, we examined the binding preferences of Tau aggregates. We found that 20 nM Tau aggregates could bind to either the pre-formed HP1α-H3K9me3-nucleosome complex or to free H3K9me3-nucleosomes, but not to free HP1α protein alone (Fig. 4g). These results indicate that Tau aggregates specifically target H3K9me3-nucleosomes rather than HP1α protein, suggesting they may disrupt HP1α chromatin localization through competitive binding. This competitive binding relationship was directly confirmed by competitive protein binding assays, which showed that Tau aggregates effectively displaced HP1α from its native binding sites on H3K9me3-modified chromatin (Fig. 4h). Furthermore, ChIP-qPCR analysis using PS19 hippocampus revealed significant enrichment of pTau at H3K9me3-enriched transposable element (TE) regions *in vivo* (Extended Data Fig. 6f). Together, these findings establish that Tau aggregates interfere with HP1α’s chromatin tethering by preferentially binding to H3K9me3-modified nucleosomes and physically displacing HP1α.

Given that Tau aggregates can bind to H3K9me3-modified nucleosomes, we investigated whether they might also interact with nucleosomes bearing other histone methylation marks. Since H3K27me3 modification also contributes to transposable element (TE) silencing^39^, we also reconstituted H3K27me3-nucleosomes (Extended Data Fig. 6g) and examined the binding capacity of Tau aggregates to H3K27me3-modified nucleosomes. Our His-pulldown assay demonstrated that Tau aggregates showed significantly weaker binding to H3K27me3-nucleosomes compared to H3K9me3-nucleosomes, while no binding was observed with unmodified nucleosomes (Extended Data Fig. 6h).

To systematically evaluate the binding preference of Tau aggregates, we employed an equimolar mixture of nucleosomes containing H3K9me3, H3K27me3, H3K36me3, and H3K4me3 modifications. Both wild-type Tau (WT-Tau) and Tau aggregates were biotin-labeled for subsequent pulldown assays. While biotin-labeled wild-type Tau (WT-Tau) showed no interaction with any nucleosome type (Fig. 4i), Tau aggregates exhibited strong preferential binding to H3K9me3-nucleosomes, substantially weaker binding to H3K27me3-nucleosomes, and negligible binding to either H3K36me3- or H3K4me3-nucleosomes (Fig. 4i). These results demonstrate that Tau aggregates selectively target H3K9me3-modified nucleosomes, suggesting their disruption of heterochromatin occurs primarily through this specific interaction.

We biochemically confirmed the direct disruptive effect of Tau aggregates on heterochromatin condensation. The condensation state of constitutive heterochromatin can be visualized through HP1α droplets formed by HP1α protein with H3K9me3-marked nucleosomal arrays^38^ (H3K9me3-NA). Notably, the HP1α droplets were completely disassembled upon addition of Tau aggregates, but not wild-type Tau (WT-Tau) (Fig. 4j).

Since the Tau aggregates used in our experiments (including those from PS19 mice) mainly contain Tau 4R repeats, while tauopathies like AD typically involve 3R/4R repeats^40^, we further examined the effects of 3R/4R Tau aggregates. We performed identical heterochromatin condensation assays using the most common AD-associated Tau isoform [Tau-3R/Tau411(P301S)]. The results demonstrated that 3R/4R Tau repeats could disrupt heterochromatin condensation as effectively as 4R Tau repeats (Fig. 4j). Notably, the decondensation of HP1α was observed notably in AD patients. The intensity of HP1α signal was attenuated especially in late-stage of AD patients (Extended Data Fig. 6i).

Thus, Tau aggregates disrupt the compactness of heterochromatin, leading to the reactivation of TEs and producing death ligand Z-RNA that triggers ZBP1-dependent necroptosis (Fig. 4k, model).

### Compromising Z-RNA-ZBP1 axis significantly attenuates AD-related phenotype in Tau-transgenic mouse model

The mechanism elucidated in this study may underlie neuronal death in tauopathies, including AD. Consistent with this model, we detected the emergence of Z-RNA (Fig. 3e) and ZBP1 (Extended Data Fig. 3i), heterochromatin erosion (Fig. 4f), and necroptotic neurons (Fig. 1g) in AD patients. Moreover, a recently published single-cell transcriptomic dataset profiling a large cohort of post-mortem AD brains^31^ revealed an inverse correlation between ZBP1 expression levels and cognitive diagnosis in AD patients (Fig. 5a). ZBP1 was significantly upregulated in excitatory neurons during disease progression from mild cognitive impairment (MCI) to dementia (Fig. 5a, left two columns), whereas its expression showed no association with cognitive status in patients with other neurological disorders (Fig. 5a, right three columns). These findings suggest that targeting the Z-RNA–ZBP1–necroptosis axis may represent a promising therapeutic strategy for AD.

**Fig. 5.**
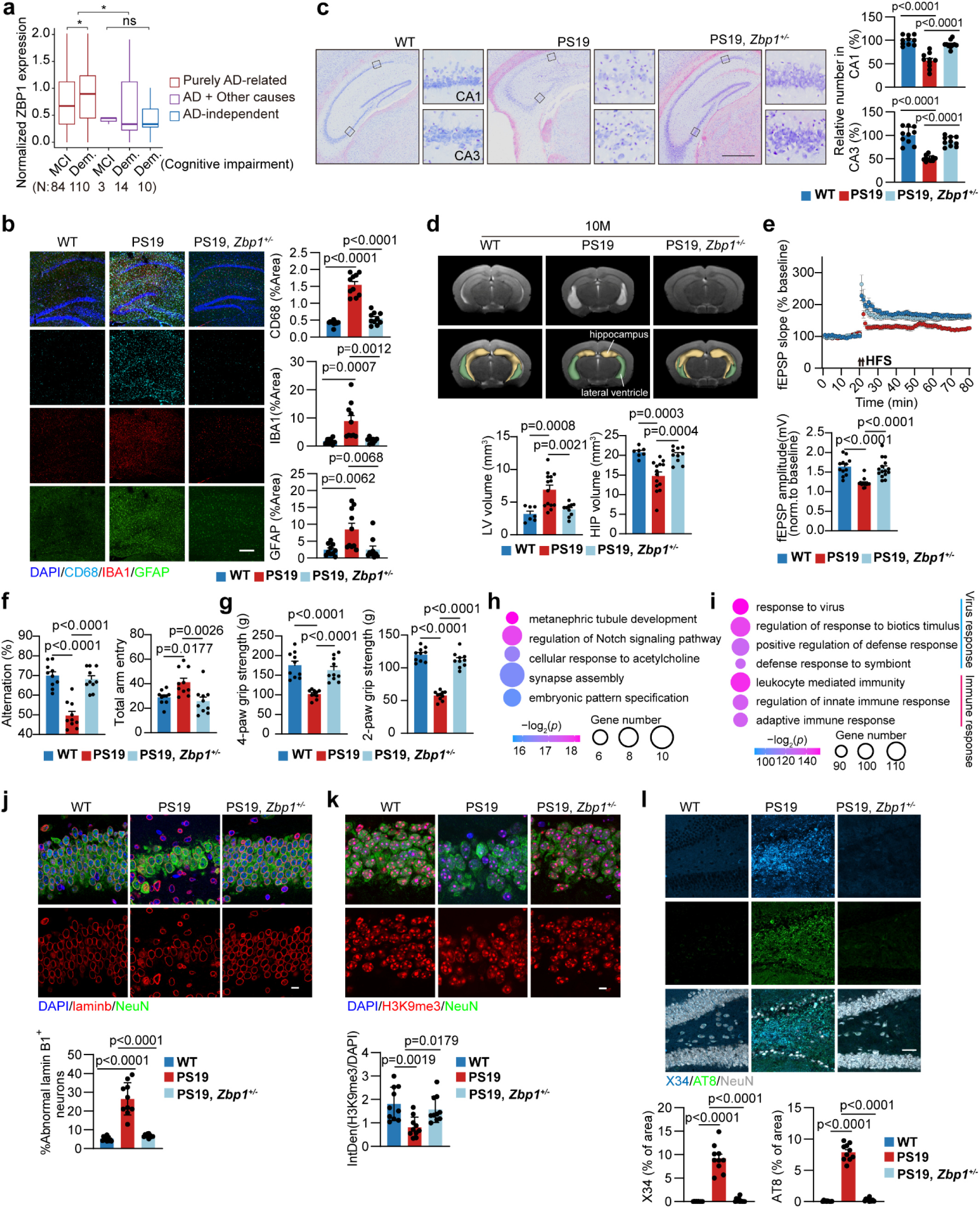
*Zbp1* haploinsufficiency rescues the neurodegenerative phonotypes in PS19 mice. **a.** Boxplot showing the quantitative comparison of ZBP1 expression between different cognitive diagnostic categories. Cognitive impairments caused by purely AD, AD coupling other causes, and AD-independent are colored in red, purple and cyan, respectively. Statistical testing was carried out by Wilcoxon rank-sum test. **b.** Representative images and quantification of area of CD68, IBA1 and GFAP on brain sections of mice with indicated genotypes. Scale bar: 300 μm. n=10 mice. **c.** Left: Nissl staining on brain sections of mice with indicated genotypes at 10-month-old. Scale bar: 700 μm. Right: quantification analysis of relative neuron number in CA1 and CA3. n=10 mice. **d.** Representative MRI images and quantification analysis of LV volume and HP volume of mice with indicated genotypes (10-month-old). WT: n=7 mice, PS19: n=14 mice, PS19, *Zbp1^+/-^*: n=10 mice. **e.** LTP recording for continuous 60 min in the hippocampal schaffer collateral region. Averaged potentiation (mean ± s.e.m.) of baseline normalized fEPSP in the indicated groups was calculated (WT: n = 6 slices from 5 mice, 12-month-old; PS19: n=7 slices from 5 mice, 10-month-old; PS19, *Zbp1^+/-^*: n=9 slices from 5 mice, 12-month-old). **f.** The cognitive function of indicated groups was evaluated by the Y-maze test for measuring total arm entries (right) and spontaneous arm alternation (left) in the indicated genotypes at 10-month-old. n=10 mice. **g.** Quantification analysis of the grip strength in mice with indicated genotypes at 10-month-old. n= 10 mice. **h.** GO enrichment analysis of downregulated genes in PS19, *Zbp1^+/-^* versus PS19 mice. **i.** GO enrichment analysis of upregulated genes in PS19, *Zbp1^+/-^* versus PS19 mice. **j.** Upper: representative images of Lamin B1 and NeuN staining in mice with indicated genotypes at 10-month-old. Scale bar: 10 μm. Lower: quantification analysis of percent of neurons with abnormal Lamin B1. n=10 mice. **k.** Upper: representative images of H3K9me3 and NeuN staining in mice with indicated genotypes at 10-month-old. Scale bar: 10 μm. Lower: quantification analysis of relative intensity of H3K9me3/DAPI. n=10 mice. **l.** Upper: representative images of X34, AT8 and NeuN staining in mice with indicated genotypes at 10-month-old. Scale bar: 50 μm. Lower: quantification analysis of area of X34 and AT8. n=10 mice. Significance between three or more groups is determined by one way ANOVA test. Data are mean ± s.e.m. * *P* ≤ 0.05; * * *P* ≤ 0.01; * * * *P* ≤ 0.001; NS, not significant.

To evaluate the therapeutic potential of necroptosis inhibition in Tau-mediated neurodegeneration, we first targeted key components of the pathway. MLKL, the terminal executor of necroptosis (Extended Data Fig. 3e), was genetically ablated. In 9-month-old PS19 mice, *Mlkl* knockout preserved brain morphology and neurological function (Fig. 1i-m). Within the necroptosis cascade, RIPK3 operates downstream of ZBP1 and upstream of MLKL (Extended Data Fig. 3e). Consistent with prior reports showing that RIPK3 inhibition attenuates neuronal death in tauopathy models (e.g., tau seeding models; see Fig.), we observed that even partial *Ripk3* deletion (heterozygous knockout) conferred neuroprotection in 10-month-old PS19 mice. This was demonstrated by restoration of neuronal density, improvement in synaptic function, reduction of neuroinflammation and rescue of cognitive deficits (Extended Data Fig. 7a-e).

We next evaluated the therapeutic potential of inhibiting Z-RNA-ZBP1 signaling in Tau pathology. Given the limited efficacy of pharmacological inhibitors, we utilized heterozygous *Zbp1* deletion (*Zbp1*^+/-^) to partially disrupt this pathway. In 10-month-old PS19 mice, *Zbp1* haploinsufficiency effectively prevented Tau aggregate-induced necroptosis (Extended Data Fig. 7f, 7g) and attenuated neuroinflammation (Fig. 5b). Remarkably, PS19,*Zbp1*^+/-^ mice exhibited preserved hippocampal neuronal counts (Fig. 5c) and maintained normal hippocampal/ventricular volumes (Fig. 5d) comparable to wild-type (WT) controls. Furthermore, synaptic dysfunction observed in PS19 mice was fully rescued in age-matched PS19, *Zbp1*^+/-^ mice by 12 months (Fig. 5e). Spinal cord analysis revealed complete absence of necroptotic neurons with significantly increased neuronal survival in PS19,*Zbp1*^+/-^ mice (Extended Data Fig. 7h, 7i). Behavioral assessments demonstrated that both spatial cognition (Fig. 5f) and motor function (grip strength, Fig. 5g) in 10-month-old PS19, *Zbp1*^+/-^ mice were indistinguishable from WT controls.

The near-complete rescue of Tau-induced neurodegeneration by ZBP1 inhibition led us to investigate the underlying molecular mechanisms. Transcriptomic analysis of hippocampi from 10-month-old PS19,*Zbp1*^+/-^ mice revealed significant upregulation of synaptic genes (Fig. 5h, Extended Data Fig. 9j), consistent with the observed synaptic functional recovery (Fig. 5e). Conversely, the most markedly downregulated genes were associated with viral infection and antiviral responses (Fig. 5i), which are typically elevated due to transposable element (TE) activation in PS19 mice. Strikingly, the expression levels of these genes in PS19,*Zbp1*^+/-^ hippocampi were restored to wild-type (WT) levels (Extended Data Fig. 7k). LncRNA-seq analysis revealed the expression level of TE was also restored to those of WT mice (Extended Data Fig. 7l), demonstrating that ZBP1 inhibition reverses TE activation.

Given that TEs are normally silenced within heterochromatin but reactivated upon heterochromatin decondensation in PS19 mice, we examined chromatin organization. PS19,*Zbp1*^+/-^ mice exhibited intact nuclear lamina architecture (Fig. 5j) and normal H3K9me3 modification patterns in heterochromatin (Fig. 5k). Since Tau aggregates disrupt heterochromatin condensation in PS19 mice (Extended Data Fig. 6c), the restored heterochromatin integrity in PS19,*Zbp1*^+/-^ mice likely reflects reduced Tau pathology. Indeed, Tau aggregates were nearly absent in hippocampi of 10-month-old PS19,*Zbp1*^+/-^ mice, contrasting with widespread aggregates in age-matched PS19 controls (Fig. 5l).

### ZBP1 Targeting Confers Long-Term Neuroprotection in Tauopathy

Although targeting MLKL or RIPK3 demonstrated neuroprotective effects in 9-10-month-old PS19 mice, long-term survival analysis revealed limitations of this approach. Notably, 3 of 20 PS19,*Mlkl*^-/-^ mice and 2 of 14 PS19,*Ripk3*^-/-^ mice died by 12 months of age (Fig. 6a). In stark contrast, upstream inhibition of ZBP1 in the death pathway provided superior protection against mortality. All 38 PS19,*Zbp1*^-/-^ and 25 PS19,*Zbp1*^+/-^ mice survived beyond 12 months (Fig. 6a), whereas only 3 of 26 age-matched PS19 control mice remained alive at this endpoint.12-month-old PS19, *Zbp1^-/-^* mice exhibited body weight (Extended Data Fig. 8a), spatial recognition ability (Extended Data Fig. 8b), limb grip (Extended Data Fig. 8c), and synaptic function (Extended Data Fig. 8d) comparable to age-matched wild-type control.

**Fig. 6.**
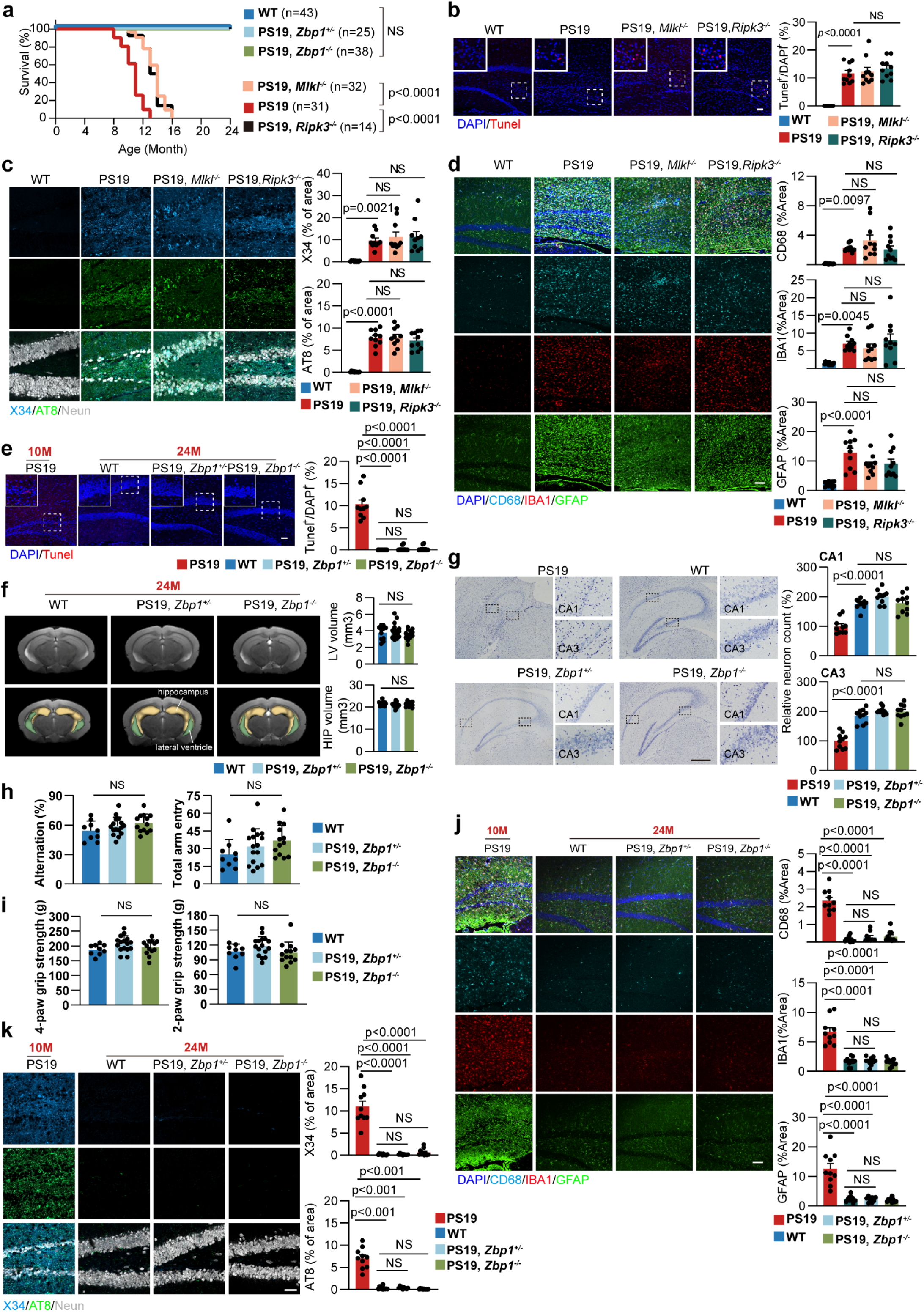
ZBP1 targeting confers long-term neuroprotection in Tauopathy. **a.** Survival curve of mice with indicated genotypes, number of mice were shown in the figure. *P* values were determined by log-rank test, 95% confidence interval of ratio. **b.** Left: representative images of Tunel staining on hippocampus sections in mice of indicated genotypes (WT: 12–13-month-old, PS19: 10-month-old, PS19, *Mlkl*^-/-^: 13–16-month-old and PS19, *Ripk3*^-/-^: 12–14-month-old). Scale bar: 50 μm. Right: quantification analysis of percent of Tunel-positive cells. n=10 mice. **c.** Left: representative images of X34, AT8 and NeuN staining in mice of indicated genotypes (WT: 12–13-month-old, PS19: 10-month-old, PS19, *Mlkl*^-/-^: 13–16-month-old and PS19, *Ripk3*^-/-^: 12–14-month-old). Scale bar: 50 μm. Right: quantification analysis of area of X34 and AT8. n=10 mice. **d.** Representative images and quantification of area of CD68, IBA1 and GFAP on brain sections of mice with indicated genotypes (WT: 12–13-month-old, PS19: 10-month-old, PS19, *Mlkl*^-/-^: 13–16-month-old and PS19, *Ripk3*^-/-^: 12–14-month-old). Scale bar: 100 μm. n=10 mice. **e.** Left: representative images of Tunel staining on hippocampus sections in mice of indicated genotypes at 24-month-old, compared with PS19 mice at 10-month-old. Scale bar: 50 μm. Right: quantification analysis of percent of Tunel-positive cells. n=10 mice. M: month. **f.** Representative MRI images and quantification analysis of LV volume and HP volume of mice with indicated genotypes at 24-month-old. WT: n=12 mice, PS19, *Zbp1^+/-^*: n=16 mice, PS19, *Zbp1^-/-^*: n=13 mice. **g.** Left: Nissl staining on brain sections of WT, PS19, *Zbp1^+/-^* and PS19, *Zbp1^-/-^* mice at 24-month-old, compared with PS19 mice at 10-month-old. Scale bar: 500 μm. Right: quantification analysis of relative neuron number in CA1 and CA3. n=10 mice. **h.** The cognitive function of indicated groups was evaluated by the Y-maze test for measuring total arm entries (right) and spontaneous arm alternation (left) in the indicated genotypes at 24-month-old. WT: n=9 mice, PS19, *Zbp1^+/-^*: n=16 mice and PS19, *Zbp1^-/-^*: n=13 mice. **i.** Quantification analysis of the grip strength in mice with indicated genotypes at 24-month-old. WT: n=9 mice, PS19, *Zbp1^+/-^*: n=16 mice, PS19, *Zbp1^-/-^*: n=13 mice. **j.** Representative images and quantification of area of CD68, IBA1 and GFAP on brain sections of WT, PS19, *Zbp1^+/-^* and PS19, *Zbp1^-/-^* mice at 24-month-old, compared with PS19 mice at 10-month-old. Scale bar: 100 μm. n=10 mice. **k.** Left: representative images of X34, AT8 and NeuN staining in mice of WT, PS19, *Zbp1^+/-^*and PS19, *Zbp1^-/-^* mice at 24-month-old, compared with PS19 mice at 10-month-old. Scale bar: 50 μm. Right: quantification of area of X34 and AT8. n=10 mice. Significance between three or more groups is determined by one way ANOVA test. Data are mean ± s.e.m. NS, not significant.

Survival curve analysis demonstrated that neither PS19,*Ripk3*^-/-^ nor PS19,*Mlkl*^-/-^ mice survived beyond 16 months of age (Fig. 6a). In end-stage PS19,*Ripk3*^-/-^ and PS19,*Mlkl*^-/-^ mice, we observed re-emergence of robust neuronal death (Fig. 6b), accumulation of prominent Tau aggregates (Fig. 6c) and exacerbation of severe neuroinflammation (Fig. 6d).These findings indicate that targeting RIPK3 or MLKL to inhibit necroptosis provides only transient neuroprotection, ultimately failing to prevent neuron death and disease progression in this Tau mouse model.

Remarkably, 2-year-old PS19,*Zbp1*^-/-^ mice and PS19,*Zbp1*^+/-^ mice maintained neurological health comparable to age-matched wild-type (WT) controls, demonstrating superior therapeutic efficacy of ZBP1 ablation compared to RIPK3 inhibition. In longitudinal survival studies revealed 100% survival (25/25) of PS19,*Zbp1*^+/-^ mice and 100% survival (13/13) of PS19,*Zbp1*^-/-^ mice, reaching 2 years without mortality or paralysis (Fig. 6a). These protected mice showed complete absence of neuronal death (Fig. 6e), prevention of brain atrophy (Fig. 6f), preservation of neuronal density (matching WT levels) (Fig. 6g), maintenance of cognitive performance (Fig. 6h, 6i), controlled neuroinflammatory responses (Fig. 6j) and significant reduction in pathogenic Tau aggregates (Fig. 6k).

### ZBP1 is not the direct upstream of Tau aggregates

In PS19,*Zbp1*^-/-^ mice, total Tau protein expression remained unchanged compared to PS19 controls (Extended Data Fig. 9a). To assess ZBP1’s role in Tau aggregation, we employed the *in vitro* Tau aggregation system in SH-SY5Y (Tau-4R) cells. IFN-induced ZBP1 overexpression in SH-SY5Y cells neither altered pathogenic Tau phosphorylation (Extended Data Fig. 9b, c) nor influenced Tau puncta formation—regardless of Tau seed induction (Extended Data Fig. 9d, e). Critically, Tau aggregates were still detectable in *Zbp1*^-/-^ primary tau-expressing neurons (Fig. 2g, 2h), demonstrating that ZBP1 ablation does not impede Tau aggregate formation.

## Discussion

Some researchers propose that neuronal death in Alzheimer’s disease (AD) represents a terminal stage of neurodegeneration, suggesting that neurons destined to die may be functionally irrecoverable even with therapeutic intervention. However, in this study, genetic ablation or knockdown of *Mlkl*, *Ripk3*, or *Zbp1*—not only prevented neuronal death but also restored neuronal activity, improved cognitive function, and reduced tau aggregate accumulation in PS19 mice (Fig. 1h–m, Fig. 5b–g, and Extended Data Fig. 7a–e). These findings indicate that neuronal death is required for Tau aggregate accumulation. Although targeting MLKL or RIPK3 conferred transient neuroprotection in 9–10-month-old PS19 mice, only ZBP1 targeting sustained this effect for up to two years. Notably, neither PS19,*Ripk3*^-/-^ nor PS19,*Mlkl*^-/-^ mice survived beyond 16 months of age (Fig. 6a). This divergence correlated with the recurrence of Tau aggregates and profound neuronal death in *Ripk3*- or *Mlkl*-deficient mice, underscoring neuronal death as a pivotal driver of Tau-mediated pathogenesis.

Tau aggregation involves both hyperphosphorylation and the presence of seeding-competent tau species. At the single-neuron level, it is difficult to reconcile how inhibition of downstream necroptotic pathways reduces upstream Tau aggregation, unless necroptosis mediators such as ZBP1 also regulate Tau expression or phosphorylation. However, our analyses show that ZBP1 does not influence these processes (Extended Data Fig. 9a–e), suggesting an alternative mechanism: neuronal death may facilitate the propagation of tau seeds between cells. Specifically, dying neurons may release tau seeds into the extracellular environment, enabling their uptake by neighboring neurons and thereby promoting tau aggregation in otherwise unaffected cells. Inhibiting necroptosis through MLKL, RIPK3, or ZBP1 could disrupt this vicious cycle of neuronal death and tau propagation, effectively shielding healthy neurons from Tau “infection”. Supporting this hypothesis, the recent phase II clinical trial reported that tau-targeting monoclonal antibodies, such as semorinemab and posidinemab, yielded cognitive benefits in AD patients^41,42^. These antibodies likely act by neutralizing extracellular tau species, which are presumably released by dying neurons—a mechanism that warrants further investigation. Collectively, these findings highlight the need to elucidate how neuronal death contributes to tau spreading.

In conclusion, our study identifies a novel mechanism underlying tau-mediated neurotoxicity and establishes ZBP1 as a potential therapeutic target. Inhibiting ZBP1 not only prevents neuronal death but also limits the propagation of pathological tau. These findings offer a promising direction for the development of disease-modifying therapies for Alzheimer’s disease and other Tauopathies.

## Acknowledgements

We thank National Health and Disease Human Brain Tissue Resource Center for providing us with brain material. We are grateful to Dr. Xin Wang (Xiamen University), Dr. Nan Liu (Zhejiang University), Dr. Guohong Li (Wuhan University) and Dr. Bing Zhu (University of Chinese Academy of Sciences) for experiment resources. The study was supported by the Zhejiang Provincial Natural Science Foundation of China (LHZSD25C070001 to W.M.), the National Natural Science Foundation of China (82530046, 82588302, 32225016 to W.M.; 82388201 to J.H. ; 32170751 to Z.-H.Y.; 82203426, 82472742 to L.S.), the National Key R&D Program of China (2024YFA1306400 and 2021YFA1101401 to W.M.; 2020YFA0803500 to J.H.), the “Pioneer” and “Leading Goose” R&D Program of Zhejiang (No.2025C02110 to W.M. and S.N.), the CAMS Innovation Fund for Medical Sciences (CIFMS) (2019-I2M-5-062 to J.H.), the Fujian Province Central to Local Science and Technology Development Special Program (2022L3079 to J.H.), the Fu-Xia-Quan Zi-Chuang District Cooperation Program (3502ZCQXT2022003 to J.H.), the Noncommunicable Chronic Diseases-National Science and Technology Major Project (2024ZD0524900 to Z.-H.Y.).

## Author contributions

Conceptualization and Supervision: W.M. and J.H.; Data curation: S.-A.W., Q.C., B.-X.Z, S.-H.G., L.L., W.S. and Z.-H.Y.; Formal analysis, X.X., J.N., J.W., Z.L., P.L., Z.-Y.C., H.-F.Y., S.Z., S.N., B.L., D.W., P.C., X.Q., Z.-H.Y., J.H. and W.M.; Methodology, S.-A.W. and Z.-H.Y.; Resource, F.D., W.C., Y.Z. and J.H.; Writing-original draft, W.M., Z.-H.Y. and Q.C.; Writing-review & editing, J.H., Q.C., Z.-H.Y. and W.M.. S.-A.W. and L.L. identified neuronal cell death was pathogenic for neurodegeneration in AD. Q.C., W.L., S.-H.G. and S.-A.W. performed most of IF/IHC and behavioral testing experiments on mouse model. B.-X.Z. performed electrophysiological experiments. S.-H.G. and B.-X.Z. performed experiments on patient samples. W.S. and P.L. performed LLPS experiments. J.N. and X.X. uncovered the negative correlation of ZBP1 level with cognitive diagnosis of AD patients. Z.-H.Y. performed cellular experiments and *in vitro* biochemical experiments. B.L., S.N. and W.C. provided pathological diagnose.

## Data and materials availability

All data are available in the main text or the supplementary materials.

## Methods

### Mice

All mice were on a C57BL/6 genetic background and housed in a constant temperature of 23℃ in a 12-hour light/dark cycle at the Laboratory Animal Center of Zhejiang University and Xiamen University. Wild-type (WT) mice were from Shanghai SLAC Laboratory Animal Co., Ltd. 5×FAD and PS19 mice were initially from The Jacksom Laboratory. *Zbp1^-/-^*, *Ripk3^-/-^*, *Mlkl^-/-^*and *Caspase 8^-/-^* mice were same as used^1,2^. *Zbp1^Za^*^1,^*^2Mut/Za^*^1,^*^2Mut^* mice were a gift from Dr. Jonathan Maelfait. PS19 mice were crossed with *Zbp1^-/-^*, *Ripk3^-/-^*, *Mlkl^-/-^*, *Zbp1^Za^*^1,^*^2Mut/Za^*^1,^*^2Mut^* or *Ripk3^-/-^, Caspase 8^-/-^* mice. Detail information of the mice strains is included in Supplementary Table 1. Both males and females were used in this research. All mouse experiments were approved by the Institutional Animal Care and Use Committed at Zhejiang University and Xiamen University.

### Human specimens

The human brain tissue was kindly provided by National Health and Disease Human Brain Tissue Resource Center. Collection of human AD specimens was approved by the local ethical committee and the institutional review board of hospitals with written informed. The detailed information of human brain specimens is shown in Supplementary Table 1.

### Hippocampal targeted injection

Ten-week-old PS19 mice were anaesthetized and mounted in a stereotaxic frame. Under aseptic conditions, 5 μg of recombined human mutant tau aggregates (2 mg/ml, Abcam, Ab246003) was carefully injected into 10-week-old PS19 mice at predetermined coordinates (from bregma, ML - 2.3 mm; AP -1.9 mm; DV -1.9 mm), compared with controls injected with PBS. After three weeks, the mice were sacrificed and the hippocampus tissues were collected for analysis.

### Tissue preparation

Mice were transcardially perfused with cold PBS. The brains and spinal cords were fixed in 4% PFA at 4℃ overnight for long fixation, or for 1h for short fixation. After being dehydrated in 20% and 30% sucrose, respectively, the samples were embedded with Tissue-Tek® O.C.T. Compound (SAKURA, 4583) and sectioned at 12 µm thickness for immunofluorescent staining. For immunohistochemistry analysis, the fixed brains were dehydrated in EtOH solutions, embedded in paraffin blocks, and sectioned at 7 μm thickness.

### Isolation and culture of cortical neurons and DRG neurons

Primary cortical neurons were isolated from mice within 2 days after birth and cultured on Poly-D-lysine coated round disks with basal Neurobasal A media supplemented with 2% B27 (Gibco, 17504-044), 1% GlutaMax (Gibco, 35050-061), 1% penicillin and streptomycin (VivaCell Biosciences, C3420-0100). After one week in culture, neurons were treated with 25 nM preformed Tau aggregates (Abcam, ab246003) to induce Tau aggregation in vitro. Control neurons were treated with recombinant human wild-type Tau441 protein (Abcam, ab84700). DRG neurons from 8-week-old PS19 mice were acquired as reported^3^. The isolated DRG neurons were cultured on Poly-D-lysine and Laminin coated round disks with basal Neurobasal A media supplemented with 2% B27, 1% GlutaMax, 20 ng/ml β-NGF (Sino Biological, 50385-MNAC), 1% penicillin and streptomycin. After culture for 3 days, 25 nM Tau aggregates were added into the culture media. To evaluate the effect of TNF on neuron death, 20 μg/ml TNF neutralizing antibody (BioXCell, BE0244) was added. After culture for another 5 days, the neurons were collected and fixed with 4% PFA for immunofluorescent staining.

### Tau seeding in biosensor cells

SY5Y tau biosensor cells stably expressing the 4R repeat domain (RD) of tau with the P301L mutation tagged with yellow fluorescent protein (YFP) were generated as previously described^4^. Cells were seeded at 2.5×10^5^ cells/mL in 500 μL of medium on Poly-L-lysine coated glass coverslips in a 24-well plate and allowed to grow overnight. The next day, 2 μg brain homogenate from Tg4510 mice was mixed with 3 μL of Lipofectamine 2000 and brought up to 50 μL in OptiMEM and allowed to sit at room temperature for 20 min. The mixture was then added to the cells and allowed to incubate at 37℃ for 6 h. The medium was changed 6 h post transfection. Tau aggregate formation was monitored using a fluorescence microscope with a 488 nm filter.

### Histology

Mouse cryosections and paraffin-sections were pre-dried at 37℃ for 2 h. For paraffin-sections, deparaffinization was performed with xylene (two washes×5 min), followed by rehydration through ethanol gradient (1 min 100% EtOH, 1 min 95% EtOH, 1 min 70% EtOH, and 1 min 50% EtOH). For immunofluorescent staining in mouse paraffin-sections, antigen retrieval was performed with citrate buffer (pH=6.0). Afterwards, the sections were washed with PBS three times, blocked with 3% BSA and 0.4% Triton X-100 at RT for 1 h, and incubated with primary antibodies including phospho-Tau (Ser202, Thr205) (AT8) (1:500, Thermo Fisher, MN1020), phospho-Tau (Ser396) (1:500, HuaBio, ET1611-68), phospho-Tau (Thr181) (1:500, CST, 12885), phospho-Tau (Thr231) (1:500, HuaBio, HA721570), phospho-Tau (Ser356) (1:500, HuaBio, HA721799), Rip3 (1:500, ProSci, 2283), pRip3 (1:500, Abcam, ab222320), pMLKL (1:500, Abcam, ab196436), Lamin b (1:500, ProteinTech, 12987-1-AP), H3K9me3 (1:2000, Abcam, ab8898) and NeuN (1:500, Oasis biofarm, OB-PGP006) overnight at 4℃. Long fixed mouse cryosections were washed with PBS three times, blocked with 3% BSA and 0.4% Triton X-100 at RT for 30 min, and then incubated with IBA1 (1:500, Oasis biofarm, OB-PGP049) and GFAP (1:2000, Oasis biofarm, OB-PGP055 and OB-MMS036). Short fixed mouse cryosections were washed with PBS, blocked with 3% BSA and 0.4% Triton X-100, followed by incubation with primary antibodies including PSD95 (1:200, Abcam, ab12093), Synap i (1:200, ImmunoWay, YT4483), AT8 (1:500), phospho-Tau (Ser396) (1:500), phospho-Tau (Thr181) (1:500), phospho-Tau (Thr231) (1:500) and phospho-Tau (Ser356) (1:500). For dsRNA or ZNA staining in mouse brain tissues, the tissues were fixed in 4% paraformaldehyde for 10 min and then subjected to proteinase K treatment (40 µg/mL) for 10-15 min at RT. When acquired, RNase A (5 mg/mL in PBS, Solarbio, 9001-99-4) or DNase I (25 U/mL in PBS, Invitrogen, AM2222) was used after proteinase K treatment at 37℃ for 1 h. The sections were incubated with primary antibodies (J2, 1:500, Sigma, MABE1134; ZNA, 1:500, Novus Biologicals, NB100-749) overnight at 4℃, washed with PBS three times, and incubated with secondary antibodies for 2 h at RT in the dark. For Thioflavin-S, Aβ (1:1000, Abcam, ab126649) and pMLKL (1:1000, Abcam, ab187091) staining, human brain sections were fixed in 4% PFA for 30 min at RT. Then sections were applied with antigen retrieval in citrate buffer and permeabilized with 0.5% Triton X-100 before being incubated with 500 uM Thioflavin-S (Sigma, T1892) for 20 min. Subsequently, after incubation in 50% ethanol for 3 min, the sections were washed with PBS for 5 min and blocked with 3% BSA at RT for 30 min. For staining including AT8 (1:500, Thermo Fisher, MN1020), H3K9me3 (1:500, Abcam, ab8898), HP1α (1:500, HuaBio, M1211-2), phospho-Tau (Thr181) (1:500, CST, 12885), Lamin b (1:500, ProteinTech, 12987-1-AP), NeuN (1:500, Oasis biofarm, OB-PGP006) in frozen human brain sections, sections were fixed in 4% PFA for 30 min at RT. Then sections were applied with antigen retrieval in citrate buffer (pH 6.0) and blocked with 3% BSA and 0.4% Triton X-100 at RT for 30 min. The sections were incubated with primary antibodies overnight at 4℃, washed with PBS three times, and incubated with secondary antibodies for 1 h at RT in the dark. For ZNA staining in frozen human brain tissues, when acquired, RNase A (5 mg/mL in PBS, Solarbio, 9001-99-4) or DNase I (250 U/mL in PBS, Invitrogen, AM2222) treatment was performed as described previously^5^ at 37℃ for 2 h before being blocked with 3% BSA and 0.4% Triton X-100 at RT for 30 min. The sections were incubated with primary antibody (ZNA, 1:500, Absolute Antibody, Ab00783-23.0) overnight at 4℃, washed with PBS three times, and incubated with secondary antibodies for 1 h at RT in the dark. Antibodies used in histology were shown in Supplementary Table 1. TUNEL staining were done with the TUNEL kit (Vazyme, A113-01) according to the manufacturer instructions. Procedure for IHC staining was referred to Wang et al^1^. DAB staining was performed with the Elivision^TM^ super HRP (Mouse/Rabbit) IHC Kit (KIT-9923, MXB biotechnology) and DAB plus kit (DAB-2032, MXB biotechnology) according to the manufacturer instructions. Nissl Staining Kit (Beyotime, C0117) was utilized for Nissl staining of cryosections according to the instructions. Images were captured by Leica SP8, Olympus FV3000 (fluorescence), and Zeiss AxioScan7 or Zeiss Axiocam 512 (bright field).

### Immunofluorescence staining

For Lamin b and ZNA staining, SY5Y cells cultured in round coverslips were fixed in 4% PFA for 10 min and washed with PBS. After being permeabilized and blocked with 0.4% Triton X-100 and 3% BSA, cells were incubated with Lamin b (1:1000, ProteinTech, 12987-1-AP) or ZNA (1:500, Absolute Antibody, Ab00783-23.0) antibody overnight at 4℃. For pMLKL, H3K9me3 and HP1α staining, SY5Y cells were fixed with 100% methanol for 10 min at -20℃, permeabilized with 0.25% TritonX-100 for 15mins at RT, and blocked with 5% BSA. Then cells were incubated with pMLKL (1:1000, Abcam, ab187091), H3K9me3 (1:2000, HuaBio, M1112-3) or HP1α (1:1000, Abcam, ab109028) antibody overnight at 4℃. For detection of Tau aggregates by Thio-S, SY5Y cells on glass coverslips were treated with Thio-S (500 µM, dissolved in 50% ethanol) for 20 min after fixation and permeabilization. Then the coverslips were washed with 50% ethanol for 3 min twice and PBS for 3 min three times, respectively. For detection of Tau aggregates by X-34, SY5Y cells on glass coverslips were incubated with X-34 (25 µM, dissolved in 40% ethanol, pH 10) for 20 min after fixation and permeabilization. Then the coverslips were rinsed with PBS for 3 min three times, incubated with the differentiation buffer (50 mM NaOH, 80% EtOH) for 2 min and rinsed with PBS for 3 min three times. The coverslips were further blocked by 3% BSA containing 0.4% Triton X-100 and stained with phospho-Tau (Ser356) (1:500, HuaBio, HA721799) antibody overnight and then visualized with Alexa Fluor-conjugated secondary antibody.

The cultured cortical neurons and DRG neurons were fixed with 4% PFA for 10 min and then washed with PBS. After being permeabilized and blocked with 0.4% Triton X-100 and 3% BSA, the neurons were incubated with β-tubulin III (1:1000, aves, TUJ), IBA1 (1:1000, Oasis biofarm, OB-PGP049), GFAP (1:2000, Oasis biofarm, OB-MMS036), AT8 (1:1000, Thermo Fisher, MN1020) and pMLKL (1:1000, Abcam, ab196436) antibodies overnight at 4℃. Cells were then washed with PBS and incubated with secondary antibodies for 1 h at RT.

For dsRNA staining, small intestinal organoids were isolated from *Villin-Cre^ERT^*^2^*;Setdb1^flox/flox^*mice^1^. After 12 hours, organoids were treated with EtOH/4-OHT for 24 hours to deleted Setdb1. At 4 days post deletion of Setdb1, organoids were fixed in 4% PFA for 15 min. After being dehydrated in 10%, 20% and 30% sucrose, respectively, organoids were embedded with OCT (SAKURA, 4583) and sectioned at 10 μm thickness for immunofluorescent staining. The sections were treated with 4% PFA for 30 min first and washed with PBS three times, then permeabilized and blocked with 0.5% Triton X-100 and 3% BSA for 30 min. Then the sections were treated with proteinase K (20 μg/ml) for 5 min at 4℃ and washed with PBS three times. Sections then were incubated with J2 (1:500, SCICONS, 10010200) antibody overnight at 4℃.

WT MEF were transfected with poly(I:C) by Lipo2000. 4.5 hours post-transfection, medium was changed, and after next 6 hours cells were fixed in 4% PFA for 30 min. After washed with PBS three times, cells were permeabilized and blocked with 0.5% Triton X-100 and 3% BSA for 30 min. Then cells were incubated with J2 (1:500, SCICONS, 10010200) antibody overnight at 4℃. Then the cells were washed with PBS, incubated with secondary antibody for 1h at RT. Images were captured by Leica SP8 and Olympus FV3000.

### Western blots

Mouse hippocampus tissues were homogenized in cold RIPA buffer (150 mM NaCl, 50 mM Tris-HCl, 0.1% SDS, 1% Triton X-100, 0.5% sodium deoxycholate, pH 7.6) supplemented with Phosphatase and Protease Inhibitor Cocktail (Abcam). After centrifugation at 12,000 rpm at 4℃ for 15 min, the supernatant of samples was collected and quantified with BCA Protein Quantification Kit (Vazyme, E112-02). Proteins of mouse hippocampus tissues were separated into cytoplasmic and nuclear fractions. Proteins were resolved by 8-12% SDS-PAGE. Antibodies utilized for Western blots were detailed in Supplementary Table 1.

### In vitro pull-down assay

Recombinant human WT Tau441 protein (Abcam, ab84700) and recombinant human Tau (mutated P301S) protein aggregate (Abcam, Active, ab246003) were purchased from Abcam. Biotin labeling for recombinant human WT or Tau aggregate was performed by using EZ-Link Sulfo-NHS-Biotin (Thermo, 21217) according to the manufactory′s instructions. Recombinant His-HP1α protein and His-tagged WT/H3K9me3-marked/H3K27me3-marked mononucleosomes were from Dr. Pilong Li’s lab (Tsinghua University). No tag H3K4me3, H3K9me3, H3K27me3 and H3K36me3-marked mononucleosomes were provided by Dr. Dong Fang’s lab (Zhejiang University). Respective recombinant proteins were mixed in the TBST buffer (20 mM Tris-HCl, pH7.6, 150 mM NaCl, 0.1% Tween-20) supplemented with Protease Inhibitor Cocktail (MCE) and then incubated at 4℃ under gentle rotation for 8 h. His-tagged proteins were pulled down with Ni-NTA agarose for 6 hours at 4℃ and biotin-tagged proteins were pulled down with Streptavidin agarose (Beyotime) for 4 hours at 4℃. For HP1α pull-down, the HP1α antibody conjugated Protein A/G agarose beads were incubated with the indicated protein mixture overnight at 4℃ under gentle rotation. Beads containing protein complexes were washed three times with TBST buffer. The immunocomplexes were eluted in non-reducing SDS sample buffer and then analyzed by western blotting.

### Y-maze test

For spontaneous alternation analysis, mice were tested in a Y-maze made from blue-painted plastic boards. The Y-shaped maze has three identical arms (50 cm long, 20 cm high and 10 cm wide) labeled as initial arm, unfamiliar arm and familiar arm. During the training period, mice were placed at the end of initial arm, allowed for freely moving between the initial arm and familiar arm for 10 min. In the test period, all arms were open and mice were recorded by a video camera (Noldus EthoVision XT) for 10 min. Total arm entries were recorded and alternation (%) was calculated as a proportion of consecutive entries in three different arms, divided by the possible alternations (total arm entries minus two).

### Grip strength test

Grip strengths of front two paws and four paws were determined with the Grip Strength Meter (Ugo Basile, Ugo 47200). According to the manufacturer’s guidance, mice were held by the tail, lowered toward the apparatus, and allowed to grab the grid using two or four paws. The mice were pulled backward horizontally and the peak tension was recorded. For each mouse, peak force was measured three times for the front two paws and four paws and averaged. A 5 min break was given for each mouse between trials.

### Electrophysiology

Mice were anesthetized with isoflurane and brains were dissected quickly. Afterwards, the brains were cut coronally at 400 μm thickness with the Vibroslice (Leica, VT 1200S) in ice-cold oxygenated (95% O_2_/5% CO_2_) artificial hypertonic glucose solution (120 mM sucrose,10 mM D-glucose, 10 mM MgSO4, 2.5 mM KCl, 0.5 mM CaCl_2_, and 64 mM NaCl, 1.25 mM NaH_2_PO4 and 26 mM NaHCO_3_). Before recording, coronal slices were recovered in oxygenated ACSF (10 mM D-glucose, 120 mM NaCl, 1.3 mM MgSO4, 3.5 mM KCl, 2.5 mM CaCl_2_, 1.25 mM NaH_2_PO4, and 26 mM NaHCO_3_) at 32℃ for 1 h and then at RT for 1 h. The Schaffer collateral inputs to the hippocampus CA1 region pathway were applied for stimulation with a bipolar stimulating electrode (FHC, Inc.). Signals were recorded by a Multi-Clamp 700B amplifier (Molecular Devices), digitized by Digidata 1550 A with glass micropipettes (1~3 MΩ) filled with ACSF. After a stable baseline recording for 20 min, two trains of 100-Hz high-frequency stimuli with an interval of 30 s were applied to induce long-term potentiation (LTP), followed by an 80-min recording. The plasticity of synaptic transmission was calculated as the initial slope (20∼80% rising phase) of field excitatory postsynaptic potentials (fEPSPs)^6^.

### Phase separation assay

HP1α protein labeled with Alexa Fluor 546 carboxylic acid (succinimidyl ester) (Thermo) and H3K9me3-marked nucleosomal arrays (NA) were produced as previously described^7^. Preparation of 384-well microscopy plates (Cellvis), including mPEGylation of silica and passivation of well with Bovine Serum Albumin (BSA) were previously described^8^. For HP1α and 12 × NA phase separation, we first diluted the NA in Phase Separation Buffer 0/500 (20 mM HEPES pH 7.4, 0 mM/500 mM NaCl), then added HP1α protein, the final NaCl concentration was 50 mM. Gently mixed and added to the well of a PEGylated and BSA passivated 384-well microscopy plate. For static imaging, recombinant Tau proteins [recombinant human WT Tau441 protein (Abcam, ab84700), recombinant human Tau (P301S) protein aggregate (Abcam, Active, ab246003) and recombinant human Tau-3R/Tau411(P301S) (from Dr. Cong Liu)] were mixed directly with HP1α and 12 × NA. For dynamic scanning, Tau proteins were added carefully after forming stable droplets of HP1α and 12 × NA. Imaging was done with a NIKON A1 microscope equipped with a 100 × oil immersion objective. NIS-Elements AR Analysis was used to analyze the images.

### Quantitative PCR with reverse transcription

Total RNA was extracted from hippocampus tissues using TRIzol reagent (Life Technologies), and reverse transcribed into cDNA. qPCR was performed with ChamQ Universal SYBR qPCR Master Mix (Vazyme) by the CFX Connect Real-Time PCR Detection System (Bio-Rad). Information for primes utilized is shown in Supplementary Table1.

### Cross-link immunoprecipitation

Cross-link immunoprecipitation (CLIP) of hippocampus tissues was referred to Wang et al^1^. Briefly, hippocampus tissues were overlaid with 1.5 mL PBS and crosslinked with 1% formaldehyde for 5 min at room temperature (RT), followed by quenching in 125 nM glycine for 5 min.The pelleted tissue was lysed in 1 ml RIPA buffer for 30 min, and then 12.5 μL RNasin Plus (Promega) was added, followed by centrifuging at 12,000 rpm for 15 min. Samples were incubated with 2 μg of anti-ZBP1 antibodies overnight at 4℃. After immunoprecipitation with Protein A/G agarose beads (Millipore), the beads were suffered from high-salt wash buffer, RIPK buffer and reverse buffer, respectively, as reported previously(16). The immunoprecipitated RNA was extracted with the TRIzol method, and purified RNA was used for RNA sequencing performed by Hangzhou Kaitai Biotechnology Co. Ltd.

### ChIP-seq and ChIP-qPCR

Standard ChIP was performed after modification of ChIP-seq performed as previously described protocol with slight modification. Approximately 1× 10^7^ cells from cortex as above was fixed in 1% formaldehyde for 10 min at room temperature (RT), followed by quenching in 125 nM glycine for 5 min. Cell nuclei were collected and lysed in lysing buffer (50 mM HEPES/KOH pH 7.6, 1 mM EDTA, 140 mM NaCl, 10% v/v Glycerol, 0.5% NP-40, 0.25% v/v Triton X-100) supplemented with protease inhibitor. Nuclei were subjected to sonication to acquire DNA fragments of 200-350 bp. One-tenth of the sonicated chromatin was reserved as input, and the remaining was used for immunoprecipitation by incubation with 2 μg H3K9me3 antibody (rabbit, Abcam, ab232324) overnight at 4°C, followed with collection using 30 μl protein A/G plus agarose beads (Millipore). After elute and purify the chromatin, ChIP-seq was performed by Hangzhou Kaitai Biotechnology Co. Ltd. For ChIP-qPCR analysis, the chromatin lysate was divided into aliquots for individual immunoprecipitation (IP) assays following sonication. These aliquots were incubated overnight at 4°C with 2 µg of the following antibodies: anti-H3K9me3, anti-pTau (rabbit, Huabio, ET1611-68) and Rabbit IgG as a negative control. The subsequent steps, including capture with Protein A/G Plus Agarose Beads, washing, elution, reversal of cross-links, and DNA purification, were performed as described in the standard protocol. For ChIP-qPCR analysis, the purified DNA (from both IP and input samples) was subjected to quantitative real-time PCR using SYBR Green Master Mix on a Bio-Rad real-time PCR system. Specific primers targeting genomic regions of interest and a negative control region were used to quantify the enrichment of H3K9me3 and pTau. The relative enrichment was calculated using the percent input method.

### Transcriptome RNA sequencing and analysis

#### Bulk RNA-seq analysis

Purified total RNA extracted from hippocampus of wild-type (WT) and PS19 mice was used for RNA-sequencing (RNA-seq) library preparation. cDNA libraries were constructed using the NEBNext Ultra RNA Library Preparation Kit and sequenced with the Illumina platform. Initial processing of FASTQ files involved filtering with Trim_Galore (v0.6.5), initially removing low-quality bases typically located at the 3’ end with quality scores < 20, followed by adapter removal. Reads shorter than 20 bp were also excluded. Processed reads were aligned to the mm39 genome assembly using Hista2, following the alignment parameters recommended by the Hista2 manual. The resulting SAM files were transformed into BAM format using Samtools. The featureCounts function from the Rsubread package (version 1.32.2) was used to quantify gene read counts. Fragments with both ends mapped exclusively were quantified at the gene level.

#### LncRNA-seq analysis

The extracted RNA underwent rRNA removal using the rRNA removal kit. Briefly, after fragmentation, first-strand cDNA synthesis occurred using random hexamer primers. Subsequently, the second-strand cDNA was synthesized, and dUTPs were substituted with dTTPs in the reaction buffer. The final library was prepared using Qubit and real-time PCR for quantification, and bioanalyzer for size distribution detection. LncRNA-seq was performed by Hangzhou Kaitai Biotechnology Co. Ltd. Newly generated sequencing data underwent quality control, wherein low-quality bases and reads less than 20 bp were removed using Trim_Galore (v0.6.5). The clean data were aligned to the mm39 reference genome using STAR. During alignment, parameters winAnchorMultimapNmax and winAnchorMultimapNmax were set to 150, in addition to default settings, to minimize multiple matching.

Transposable element (TE) expression was quantified using RepeatMasker annotations (BED format) from the UCSC Genome Browser. RNA-seq alignments were converted to BED using bedtools v2.30.0 bamtobed, and overlapping reads were identified with bedtools intersect. For each repeat element, uniquely mapped reads were counted, and only elements with ≥5 reads were retained to reduce low-abundance noise. Counts were normalized to reads per million mapped reads (RPM) to account for sequencing depth differences. RNAs with Log2 Fold Change (LFC) > 2 and Benjamini-Hochberg-corrected p-value < 0.05, relative to their WT levels, were categorized as enriched. Heatmaps were generated using the ComplexHeatmap package (v2.16.0).

#### RIP-seq analysis analysis

For RIP-seq analysis, the adapter sequences of raw reads were initially removed using cutadapt v4.0. Subsequently, the processed reads were aligned to the mouse genome mm10 using Hisat2 v2.2.1 with default parameters. Peak calling was done using macs2 v2.2.7.1 with these key parameters: -f BED – nolambda – nomodel – extsize 50. To obtain reliable peaks in the RIP-seq data, the following filtering criteria were applied: (1) maximum count in the IP group > 10; (2) log2 fold change of peak RPM (IP vs. input) > 0;(3) P-value < 0.05 (two-sided t-test). Peak distribution across genomic regions was calculated using HOMER v4.11 annotatePeaks.pl script from HOMER v4.11. Differential peaks were defined as peaks with P value less than 0.05 (two-sided t-test) and fold change larger than 2. The following Pathways and GO analysis were annotated in the KEGG (Kyoto Encyclopedia of Genes and Genomes) database phyper and shown by ggplot2 (https://ggplot2.tidyverse.org/).

#### ChIP-seq analysis

We firstly performed the quality and adapter trimming of the raw reads using the TrimGalore program (https://github.com/FelixKrueger/TrimGalore). Clean reads were then aligned to the mouse mm10 reference genome via Bowtie v.1.2.21. SAMtools was applied for the BAM file conversion and duplicates removal. Peak calling was conducted with MACS2 v2.2.7.13 using following key parameters: -f BAMPE --keep-dup all --nomodel --extsize 200. We obtained the peaks based on P-value < 0.05 (two-sided t-test). Differential peaks for the PS19 vs. WT comparisons were selected using the threshold of P-value (two-sided t-test) < 0.05 and log2 fold change in reads per million of mapped reads (RPM) < -1.

### ZBP1 expression analysis in single-cell transcriptomic data

We employed a publicly-available single-cell transcriptomic dataset that profiled a large number of AD post-mortem brains, and interrogated the expression changes of *ZBP1* in different stages of AD progression^9^. We generated a pseudobulk gene expression matrix based on the cell annotation reported by the original paper, and carried out z-score normalization to facilitate differential expression analysis. To mitigate the effect caused by outlier values in the single-cell data, we removed the individuals with the highest 10% and lowest 10% ZBP1 expression. The ZBP1 expression was compared between different cogdx scores^10^, which partitioned cognitive diagnostics based on the relevance to AD. Statistical testing between groups was carried out using Wilcoxon rank-sum test. All source data for LncRNA-seq, bulk RNA-seq and RIP-seq have been deposited at Gene Expression Omnibus under accession code GSE248070 and are publicly available as of the date of publication.

### MRI to evaluate brain atrophy

We conducted MRI scanning experiments on PS19 (9M n=11, 10M n=14); PS19,DKO (9M n=8); PS19, *Mlkl^-/-^* (9M n=5, 11M n=6); PS19, *Zbp1^+/-^* (10M n=10) and WT (10M n=7) mice using a Bruker 9.4T MRI system (BioSpec 94/30 USR) with an internal diameter of 72 mm. Mice were anesthetized with isoflurane (RWD, R500IE) and T2-weighted 2D Fast Spin-Echo Imaging were performed with rapid-acquisition-with-relaxation-enhancement(RARE). The scanning parameters: FOV = 20 x 20 mm^2^, Matrix Size = 200 x 200, TE = 33ms, TR =3000ms, RARE factor =8, Slice Thickness =0.5mm, Slice Numbers =20, number of averages = 4. 3D Slicer was used for volume analysis. Outline the hippocampus and lateral ventricles using “segment editor”, and 3D Slicer will automatically calculate the volume of hippocampus and lateral ventricles.

### Statistical analysis

ImageJ was used for quantification of immunohistochemistry and immunoblot data. The number of replicates or experiments are indicated in the individual figure legends. Data were shown in graphs as mean±s.e.m. Unless specifically indicated, all values were from at least three independent biological replicates. The two-tailed unpaired method (Student’s *t*-test) for two independent groups, and log-rank test for survival curves were utilized for significance analysis, set at a 95% confidence interval. Significance between three or more groups is determined by one way ANOVA test. Statistical analysis was performed with GraphPad Prism 8 and Microsoft 2017 software.

### Data availability statement

The authors declare that the data supporting the findings of this study are available within the paper. All source data for LncRNA-seq (GSE306753), bulk RNA-seq (GSE306761), ChIP-seq (GSE306751) and RIP-seq (GSE306739) will be deposited at Gene Expression Omnibus and are publicly available as of the date of publication.

---

**Extended Data Fig. 1.**
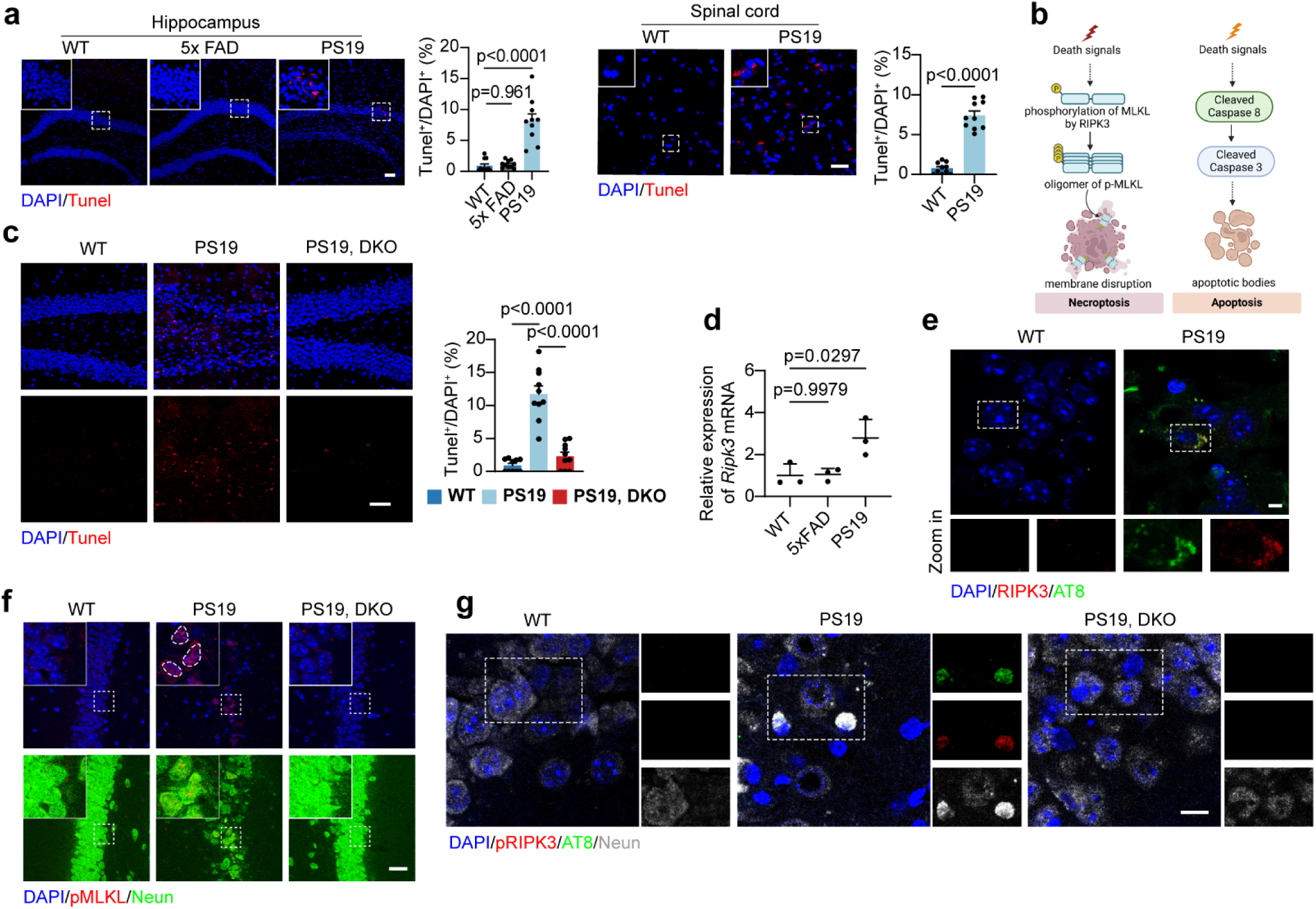
Neuron necroptosis is essential for Tau-induced neurodegeneration. **a.** Representative images of Tunel staining and quantification of Tunel/DAPI on hippocampus sections of mice with indicated genotypes and on spinal cord (SC) section of PS19 mice (WT: 9-month-old, 5xFAD: 7-month-old and PS19: 9-month-old). Scale bar: 50 μm. n=10 mice. Selected areas are shown at higher magnification. **b.** Schematic of classical necroptosis and apoptosis pathway. **c.** Representative images of Tunel staining and quantification of Tunel/DAPI on hippocampus sections from mice with indicated genotypes (9-month-old). Scale bar: 50 μm. n=10 mice. **d.** RT–qPCR analysis of relative *Ripk3* mRNA expression level in hippocampus of mice with indicated genotypes. n=3 mice. **e.** RIPK3 and p-Tau (AT8) staining on brain sections of mice with indicated genotypes. Selected areas are shown at higher magnification. Scale bar: 5 μm. **f.** Representative images of pMLKL co-stained with NeuN on brain sections of mice with indicated genotypes at 9-month-old. Selected areas are shown magnified to the left of each image. Scale bar: 20 μm. **g.** Representative images of pRIPK3 co-stained with p-Tau (AT8) and NeuN on brain sections of mice with indicated genotypes at 9-month-old. Selected areas are shown magnified to the right of each image. Scale bar: 10 μm. Significance between two groups is determined by unpaired *t*-test. Significance between three groups is determined by one way ANOVA test. Data are mean ± s.e.m.

**Extended Data Fig. 2.**
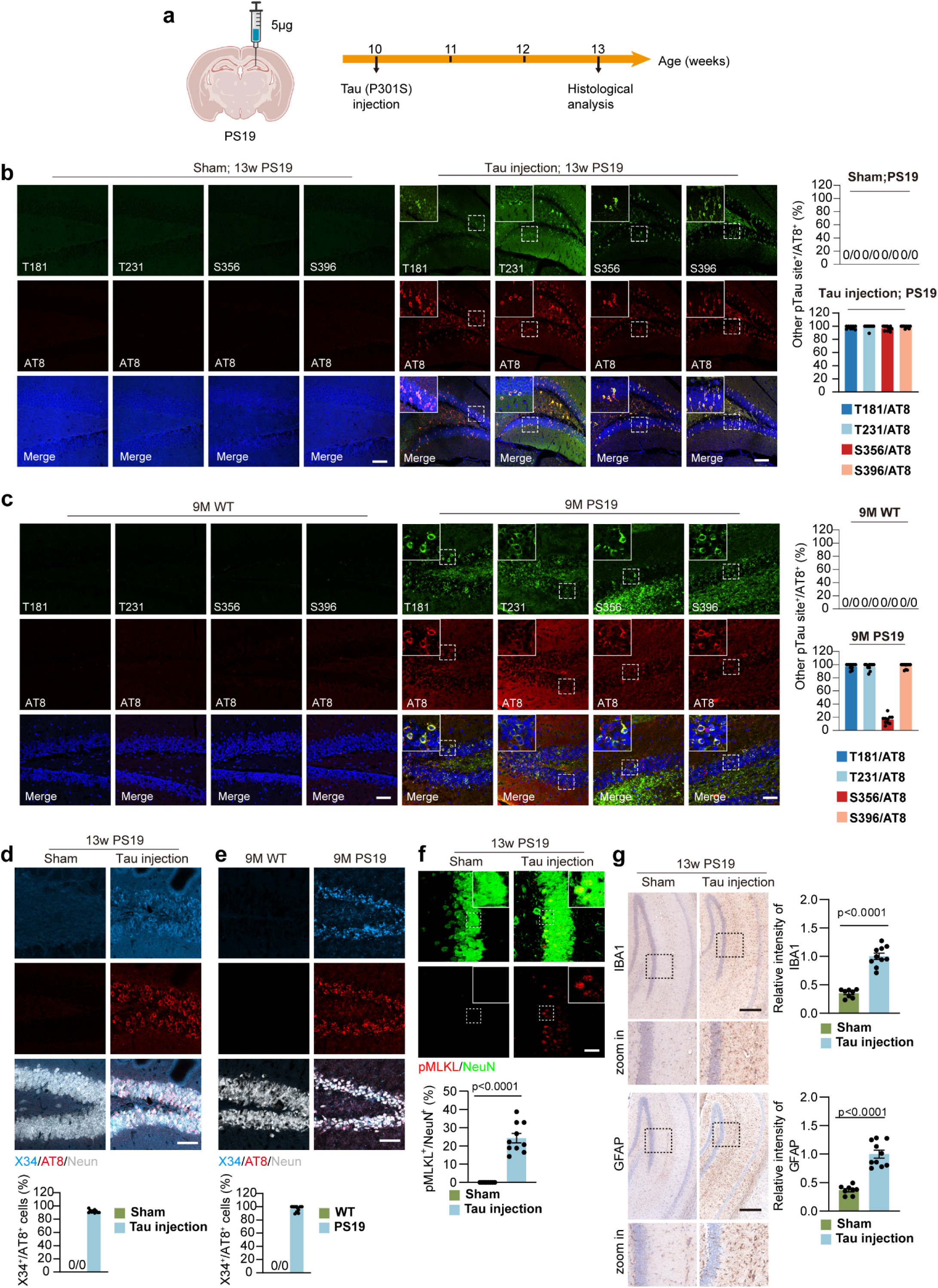
Characterization of pathological Tau in mouse model. **a.** Schematic drawing of tau acceleration model. **b.** Left: representative images of staining of various phospho-Tau epitope on brain sections from PS19 mice treated with mutant Tau. Selected areas are shown magnified to the left of each image. Scale bar: 100 μm. Right: quantification analysis of the percent of each phospho-Tau epitope in AT8-positive cells. n=10 mice. 13w: 13-week-old. **c.** Left: representative images of staining of various phospho-Tau epitope on brain sections from 9-month-old (9M) PS19 mice. Selected areas are shown magnified to the left of each image. Scale bar: 50 μm. Right: quantification analysis of the percent of each phospho-Tau epitope in AT8-positive cells. n=10 mice. **d.** Upper: representative images of X34 and AT8 staining on brain sections from PS19 mice treated with mutant Tau. Scale bar: 50 μm. Lower: quantification analysis of the percent of X34 in AT8-positive cells. n=10 mice. **e.** Upper: representative images of X34 and AT8 staining on brain sections from 9-month-old PS19 mice. Scale bar: 50 μm. Lower: Quantification analysis of the percent of X34 in AT8-positive cells. n=10 mice. **f.** Upper: representative images of pMLKL and NeuN staining on brain sections from PS19 mice treated with mutant Tau. Scale bar: 10 μm. Lower: quantification analysis of the percent of pMLKL-positive neurons. n=10 mice. **g.** Left: representative images of IBA1 and GFAP staining on brain sections from PS19 mice treated with mutant Tau. Scale bar: 300 μm. Selected areas are shown at higher magnification. Right: quantification of relative intensity of IBA1 and GFAP. Sham PS19: n=8 mice, Tau injected PS19: n=10 mice. Significance between two groups is determined by unpaired *t*-test. Data are mean ± s.e.m.

**Extended Data Fig. 3.**
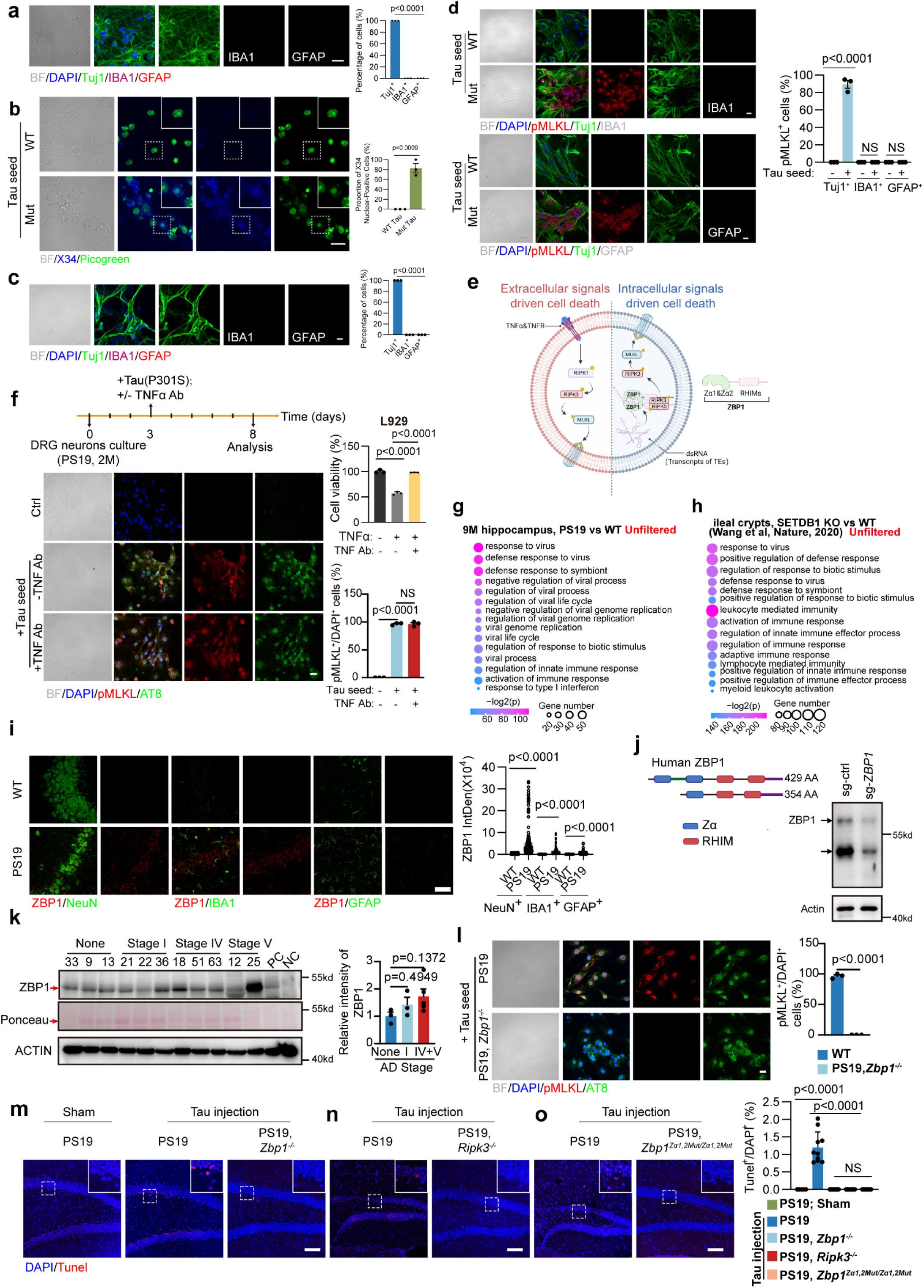
Tau aggregate induces ZBP1-dependent neuronal necroptosis. **a.** Left: representative images of Tuj1, IBA1 and GFAP staining and bright field of primary cortex neurons cultured *in vitro*. Scale bar: 20 μm. Right: quantification analysis of percent of Tuj1, IBA1 and GFAP positive cells. n=3 independent experiments. **b.** Left: representative images of X34, PicoGreen staining and bright field of primary cortex neurons cultured *in vitro*. Scale bar: 20 µm. Right: quantification analysis of the percentage of X34 nuclear positive cells. n=3 independent experiments. **c.** Left: representative images of Tuj1, IBA1 and GFAP staining and bright filed of primary DRG neurons cultured *in vitro*. Scale bar: 20 μm. Right: quantification analysis of percent of Tuj1, IBA1 and GFAP positive cells. n=3 independent experiments. **d.** Left: representative images of pMLKL, Tuj1, IBA1 and GFAP staining and bright field of primary PS19 DRG neurons treated with recombinant WT or mutant Tau protein. Scale bar: 20 μm. Right: quantification analysis of pMLKL-positive cells. n=3 independent experiments. **e.** Schematic drawing of extracellular signal-driven cell death (left) and intracellular signal-driven cell death (right). **f.** Left upper: schematic drawing of PS19 DRG neuron culture with treatment of TNFα antibody. Left lower: representative images of pMLKL and AT8 staining and bright filed of primary DRG neurons treated recombinant WT or mutant Tau protein with or without TNF antibody. Scale bar: 20 μm. Right upper: the efficiency of TNF antibody was tested by examining the cell viability of L929 treated with TNFα with or without TNF antibody. Right lower: quantification of pMLKL-positive cells treated with TNF antibody or not was performed. n=3 independent experiments. **g.** Global transcriptome analysis of hippocampus from 9-month-old mice (PS19 versus WT). **h.** Global transcriptome analysis of ileal crypts (SETDB1 KO versus WT) from published work (Wang et al, Nature, 2020). **i.** Upper: representative images of ZBP1, NeuN, IBA1 and GFAP staining on brain sections of WT and PS19 mice (9-month-old). Scale bar: 50 μm. Lower: quantification of intensity of ZBP1 in NeuN^+^, IBA1^+^ and GFAP^+^ cells. **j.** Left: Schematic drawing of two isoforms of human ZBP1. Right: Immunoblotting analysis of ZBP1 protein level in H929 cells transduced with Lenti-CRISPR/Cas9-gRNA construct targets targeting *ZBP1* or Ctrl. **k.** Immunoblotting and quantification of ZBP1 protein level in the hippocampal protein samples of AD patients at various stage as indicated. Sample from IFN-treated SH-SY5Y cells was used as a positive control (PC). **l.** Left: representative images of pMLKL (red) and AT8 (green) staining and bright filed of primary DRG neurons from PS19 and PS19, *Zbp1*^-/-^ mice cultured *in vitro*. Scale bar: 20 μm. Right: quantification analysis of percent of pMLKL-positive cells. n=3 independent experiments. **m-o.** Representative images and quantification analysis of percent of neuronal death (Tunel-positive) in Tau acceleration model with indicated genotypes (Sham PS19: n=5 mice, PS19: n= 10 mice; PS19, *Zbp1^-/-^*: n=7 mice; PS19, *Ripk3^-/-^*: n=6 mice; PS19, *Zbp1^Za^*^1,^*^2Mut/Za^*^1,^*^2Mut^*: n=6 mice). Selected areas are shown magnified to the right of each image. Scale bar: 20 μm. Significance between two groups is determined by unpaired *t*-test. Significance between three or more groups is determined by one way ANOVA test. Data are mean ± s.e.m. NS, not significant.

**Extended Data Fig. 4.**
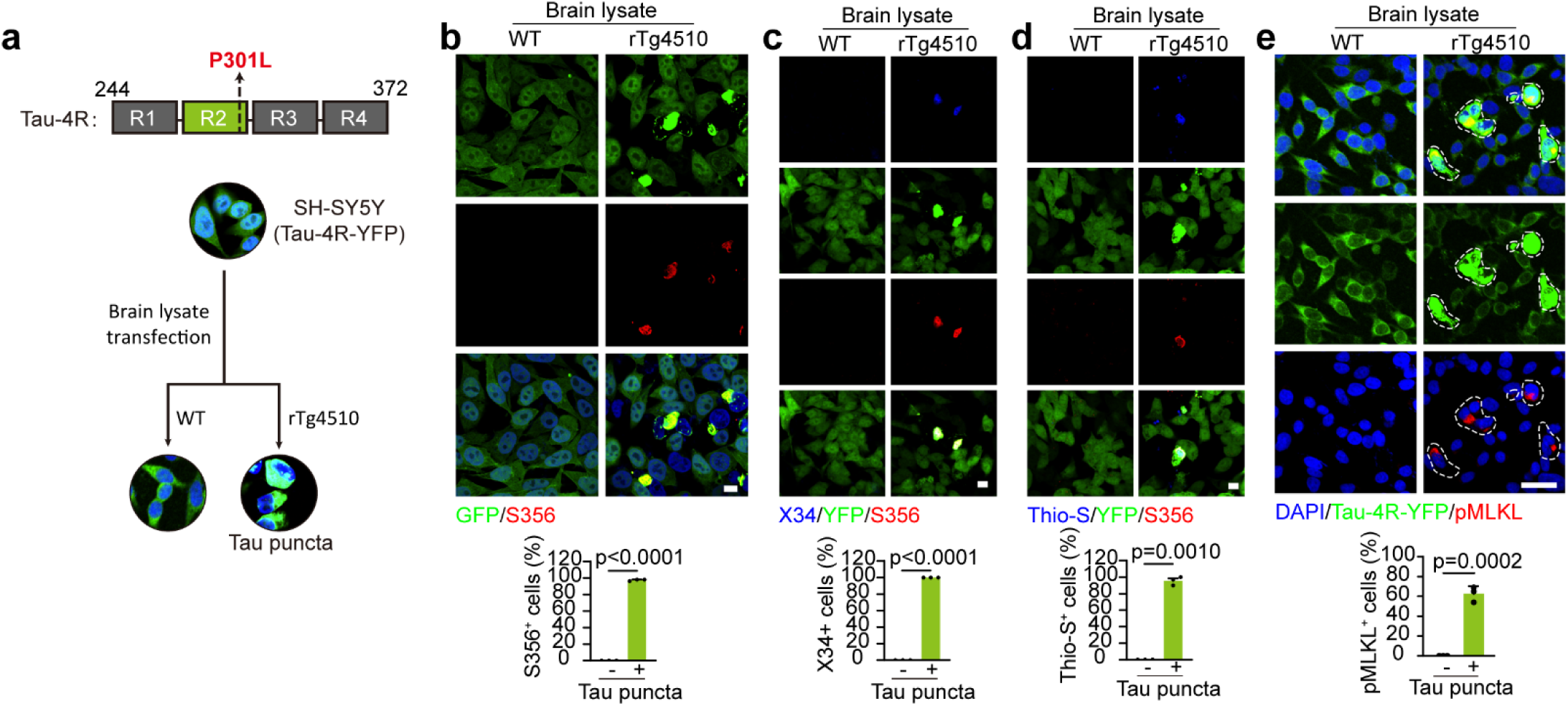
Characterization of pathological Tau in SH-SY5Y (Tau-4R-YFP) cells. **a.** Schematic drawing of the tau seeding in SH-SY5Y-Tau 4R cells (SH-SY5Y stably expressed with Tau-4R-YFP). **b.** SH-SY5Y-Tau 4R cells were treated with brain lysates from rTg4510 or WT mice. Upper: representative images of phospho-Tau (S356) signal (red) in tau puncta (green) containing cells. Scale bar: 20 μm. Lower: quantification analysis of percent of S356-positive cells in Tau puncta containing cells. n=3 independent experiments. **c.** SH-SY5Y-Tau 4R cells were treated with brain lysates from rTg4510 or WT mice. Upper: representative images of phospho-Tau (S356) signal (red) with X34 in tau puncta (green) containing cells. Scale bar: 20 μm. Lower: quantification analysis of percent of X34-positive cells in Tau puncta containing cells. n=3 independent experiments. **d.** SH-SY5Y-Tau 4R cells were treated with brain lysates from rTg4510 or WT mice. Upper: representative images of phospho-Tau (S356) signal (red) with Thio-S on tau puncta (green) containing cells. Scale bar: 20 μm. Lower: quantification analysis of percent of Thios-positive cells in Tau puncta containing cells. n=3 independent experiments. **e.** SH-SY5Y-Tau 4R cells were treated with brain lysates from rTg4510 or WT mice. Upper: representative images of pMLKL signal (red) on tau puncta (green) containing cells. Scale bar: 20 μm. Lower: quantification analysis of percent of pMLKL-positive cells in Tau puncta containing cells. n=3 independent experiments. Significance between two groups is determined by unpaired *t*-test. Data are mean ± s.e.m.

**Extended Data Fig. 5.**
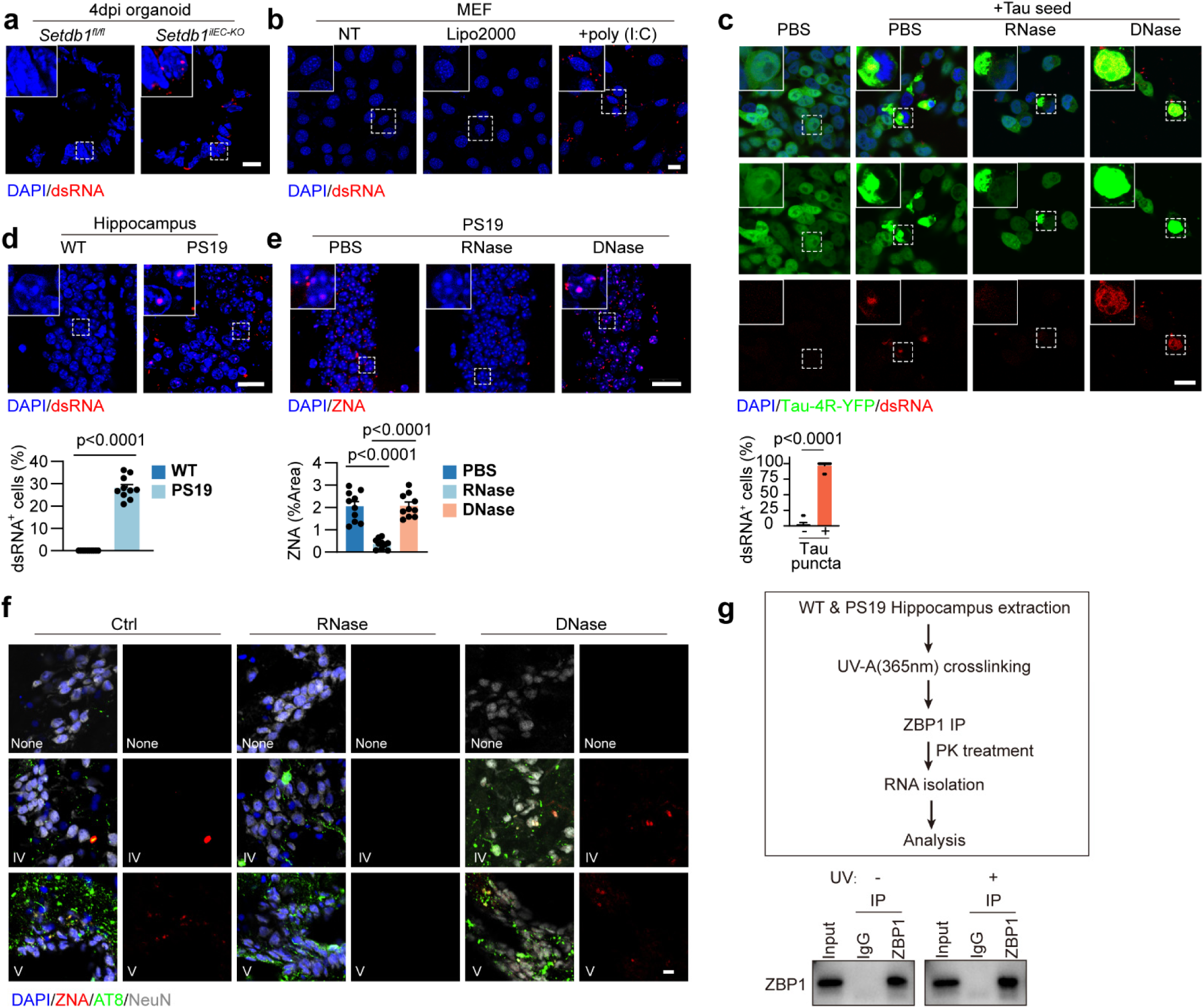
Tau aggregates produced Z-RNAs from reactivated TEs. **a.** dsRNA staining of intestinal organoid from mice with indicated genotypes. Scale bar: 20 μm. **b.** dsRNA staining of MEF cell transfected with poly(I:C). Scale bar: 20 μm. **c.** Upper: representative images of dsRNA staining in SY5Y-Tau 4Rcells treated with Tau seeds or control. The dsRNA signal was further confirmed with RNase or DNase treatment. Lower: quantification of dsRNA signal co-localized with Tau puncta. n=3 independent experiments. **d.** Upper: dsRNA staining on brain sections from mice with indicated genotypes at 9-month-old. Scale bar: 20 μm. Lower: quantification of dsRNA-positive cells. **e.** Upper: representative images of ZNA staining on brain sections from PS19 mice at 9-month-old with RNase, DNase or PBS treatment. Scale bar: 20 μm. Lower: quantification of area of ZNA on brain sections. **f.** Representative images of ZNA (with RNase or DNase treatment) and AT8 staining on brain sections from AD patients at indicated stage. Scale bar: 20 μm. **g.** Upper: working flow of RIP-seq. Lower: Immunoblotting analysis of ZBP1-IP efficiency as indicated. Significance between two groups is determined by unpaired *t*-test. Significance between three or more groups is determined by one way ANOVA test. Data are mean ± s.e.m.

**Extended Data Fig. 6.**
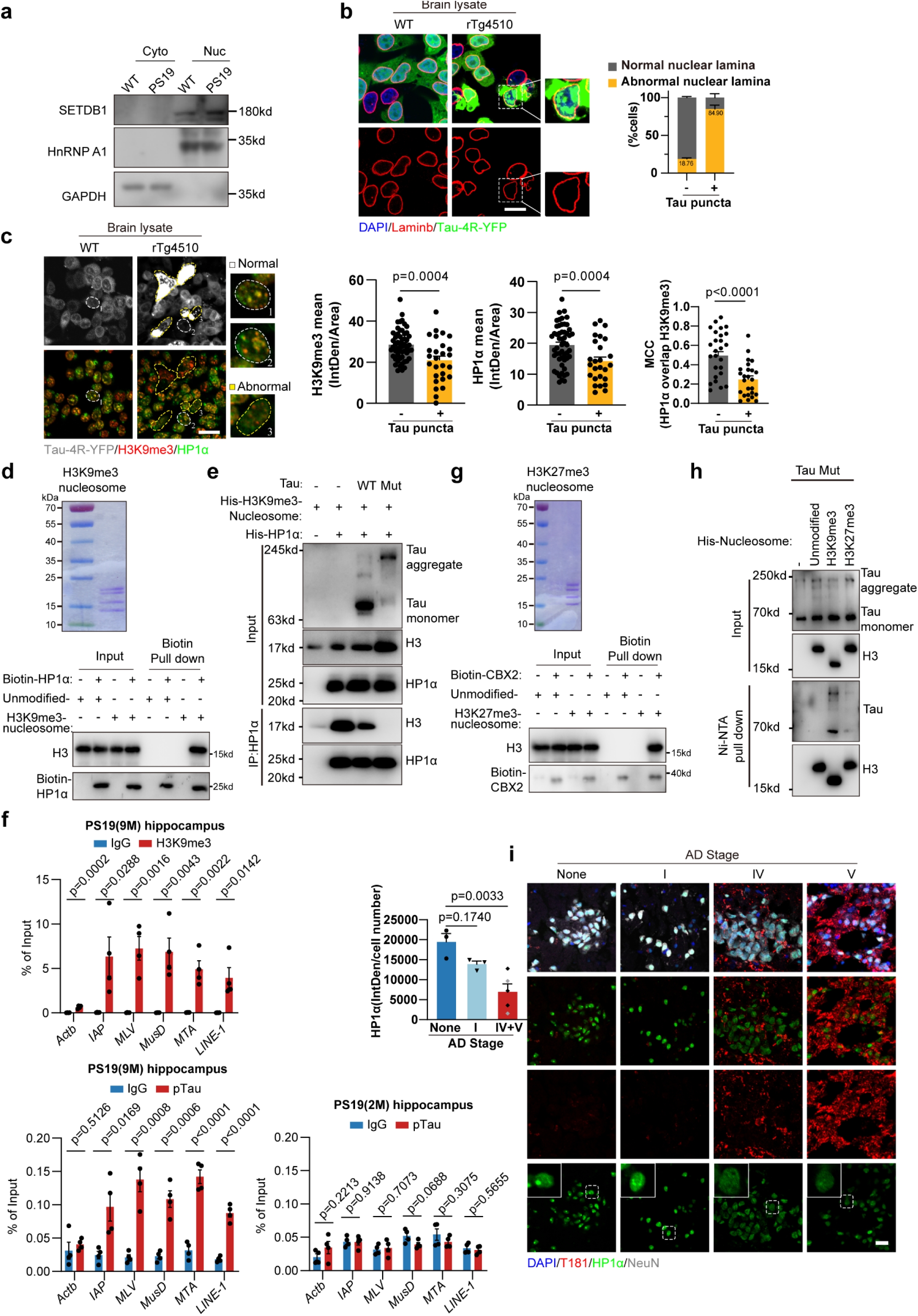
Pathogenic Tau deforms heterochromatin condensation. **a.** Immunoblotting analysis of SETDB1 in hippocampus of WT and PS19 mice (9-month-old). **b.** Left: representative images of Lamin B1 staining on SH-SY5Y (Tau-4R-YFP) cells treated with Tau seed from rTg4510 or WT mice. Scale bar: 20 μm. Selected areas are shown magnified to the right. Right: quantification of percent of cells with normal/abnormal nuclear lamina in Tau puncta-positive SH-SY5Y (Tau-4R-YFP) cells. n=3 independent experiments. **c.** Left: representative images of H3K9me3 (red) and HP1α (green) staining on SH-SY5Y (Tau-4R-YFP) cells treated with Tau seed from rTg4510 or WT mice. Scale bar: 20 μm. Selected areas are shown magnified to the right. Middle: quantification of percent of cells with HP1α and H3K9me3 overlap signal in Tau puncta-positive Tau-4R-YFP cells was performed. Right: quantification of the intensity of H3K9me3 and HP1α signal in tau puncta containing cells. n=3 independent experiments. **d.** (Top) Quality assessment by Coomassie blue staining of the recombinant H3K9me3-nucleosomes used in the pull-down assay. (Bottom) Pull-down assay demonstrates a direct interaction between H3K9me3-modified nucleosomes and the reader protein HP1α. **e.** WT or mutant tau protein was incubated with His-tagged HP1α and H3K9me3-marked nucleosome as indicated. The inputs and anti-HP1α immunoprecipitates were analyzed by immunoblotting as indicated. **f.** ChIP-qPCR analysis of (Top) H3K9me3 and (Bottom) pTau occupancy at transposable element (TE) loci in the hippocampus of 9-month-old (9M) PS19 mice. The specificity of pTau antibody was validated in 2-month-old (2M) PS19 mice, which show no detectable pTau signal at this age. Data represent the mean percentage of input (± s.e.m). The *Actb* gene, which is not expected to bind these factors, was included as a negative control. **g.** (Top) Quality assessment by Coomassie blue staining of the recombinant H3K27me3-nucleosomes. (Bottom) Pull-down assay confirms the specific binding of CBX2 to H3K27me3-modified nucleosomes. **h.** Recombinant mutant tau protein (20 nM) was incubated with His-tagged unmodified, H3K9me3-marked or H3K27me3-marked nucleosome respectively. Inputs and Ni-NTA pull-down samples were analyzed by immunoblotting as indicated. Note: the histone 3 in the unmodified and H3K27me3-marked nucleosome included an additional Flag tag. **i.** Left: representative images of p-Tau (T181), HP1αand NeuN staining on hippocampus sections of AD patients at various stage as indicated. Scale bar: 20 μm. Selected areas are shown at higher magnification. Right: quantification analysis of intensity of HP1α signal on hippocampus of AD patients. Non-AD: n=3, Stage I: n=3, Stage IV+V: n=5. Significance between two groups is determined by unpaired *t*-test. Significance between three or more groups is determined by one way ANOVA test. Data are mean ± s.e.m.

**Extended Data Fig. 7.**
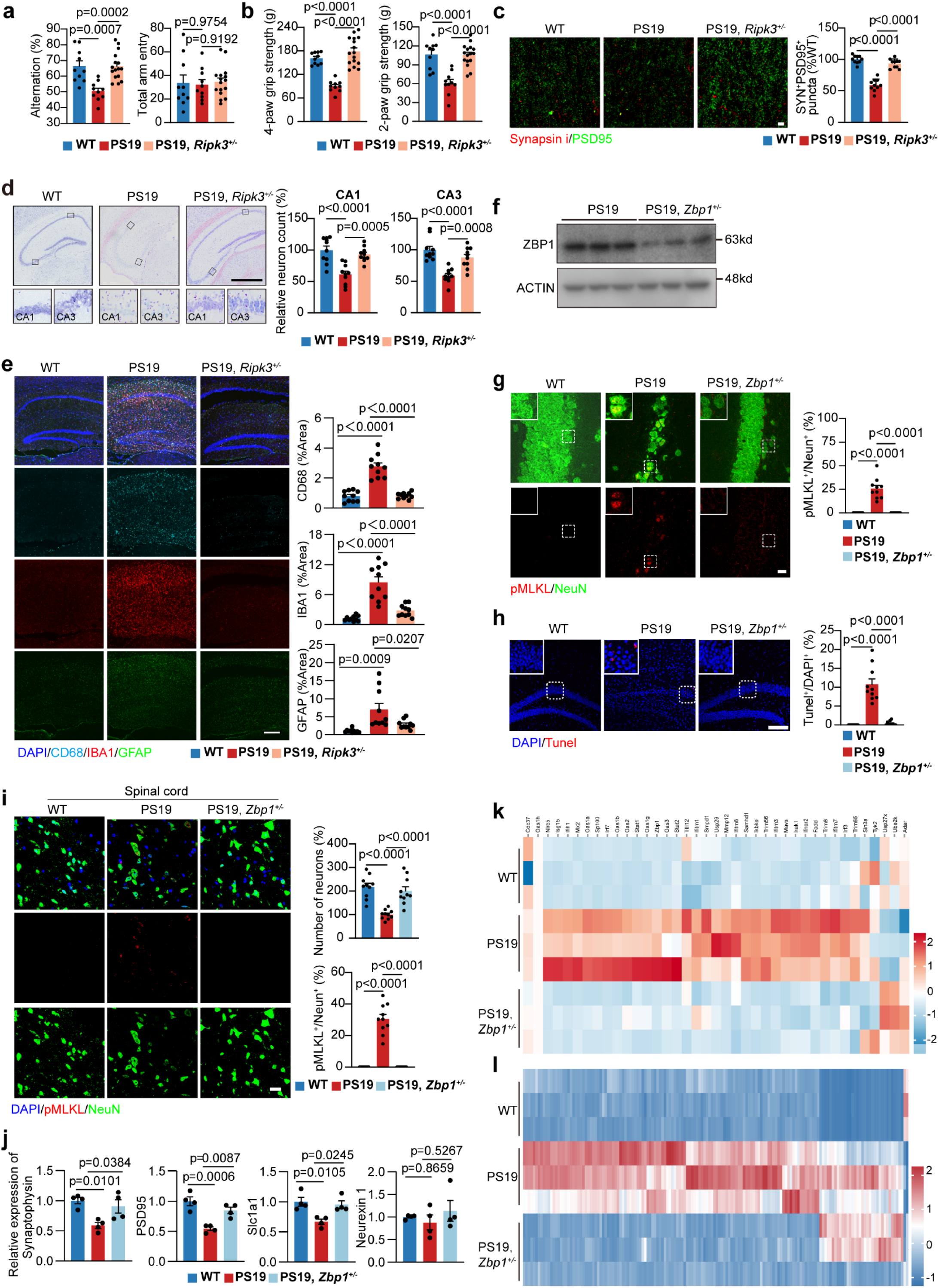
*Zbp1* haploinsufficiency effectively rescues the neurodegenerative phenotypes in PS19 mice. **a.** The cognitive function of indicated groups was evaluated by the Y-maze test for measuring spontaneous arm alternation (left) and total arm entries (right) in mice with the indicated genotypes at 10-month-old. WT: n=10 mice, PS19: n=10 mice, PS19, *Ripk3*^+/-^: n=16 mice. **b.** Quantification analysis of the grip strength in mice with indicated genotypes at 10-month-old. WT: n=10 mice, PS19: n=10 mice and PS19, *Ripk3*^+/-^: n=16 mice. **c.** Left: super-resolution images of Synapsin i (red) and PSD95 (green) immunoreactive puncta in the hippocampus of mice with the indicated genotypes at 10-month-old. Scale bar: 5 μm. Right: quantification of SYN^+^PSD95^+^ puncta within groups. n=10 mice. **d.** Left: Nissl staining on brain sections of mice with indicated genotypes at 10-month-old. Scale bar: 700 μm. Lower: quantification analysis of relative neuron number in CA1 and CA3. n=10 mice. **e.** Representative images and quantification of area of CD68, IBA1 and GFAP on brain sections of indicated genotypes at 10-month-old. Scale bar: 300 μm. n=10 mice. **f.** Immunoblotting of ZBP1 protein level in hippocampus of PS19 and PS19, *Zbp1^+/-^* mice at 10-month-old. n=3 mice. **g.** Left: representative images of pMLKL /NeuN staining on hippocampus of mice with indicated genotypes at 10-month-old. Scale bar:10 μm. Right: quantification analysis percent of pMLKL-positive neurons. n=10 mice. **h.** Left: representative images of Tunel staining on hippocampus of indicated genotypes at 10-month-old. Scale bar: 50 μm. Right: quantification analysis of percent of Tunel-positive cells. n=10 mice. **i.** Left: representative images of pMLKL /NeuN staining on spinal cord of mice with indicated genotypes at 10-month-old. Scale bar: 20 μm. Right: quantification analysis of percent of pMLKL-positive neurons and neuron numbers. n=10 mice. **j.** Relative mRNA expression of *Synaptophysin*, *PSD95*, *Slc1a1* and *Neurexin 1* in hippocampus of mice with indicated genotypes at 10-month-old. n=4 mice. **k.** Heat map analysis of type I interferon-related genes in the hippocampus from mice with indicated genotypes. n=3 mice **l.** Heat map analysis of TEs in the hippocampus from mice with indicated genotypes. n=3 mice Significance between three or more groups is determined by one way ANOVA test. Data are mean ± s.e.m.

**Extended Data Fig. 8.**
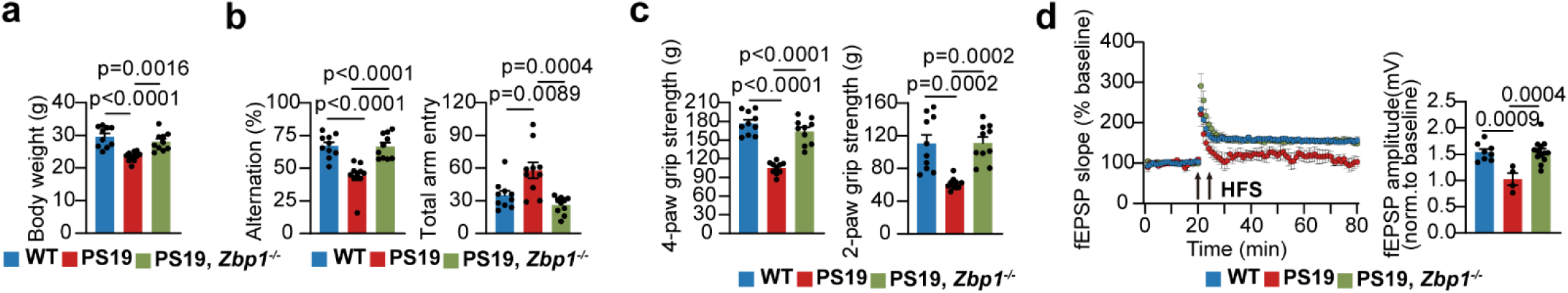
*Zbp1* deletion rescues the neurodegenerative phenotypes in PS19 mice. **a.** Quantification analysis of body weight of PS19, *Zbp1*^-/-^ and WT mice at 12-month-old, and PS19 mice at 10-month-old. n =10 mice. **b.** The cognitive function evaluated by the Y-maze test for measuring total arm entries (right) and spontaneous arm alternation (left) in PS19, *Zbp1^-/-^* and WT mice at 12-month-old, and PS19 mice at 10-month-old. n =10 mice. **c.** Quantification analysis of the grip strength in PS19, *Zbp1^-/-^* and WT mice at 12-month-old, and PS19 mice at 10-month-old. n =10 mice. **d.** LTP recording for continuous 60 min in the hippocampal Schaffer collateral region of mice with indicated genotypes at 12-month-old. Averaged potentiation (mean ± s.e.m.) of baseline normalized fEPSP in the indicated groups was calculated WT: n = 8 slices from 3 mice, PS19: n= 4 slices from 3 mice, PS19, *Zbp1^-/-^*: n=13 slices from 3 mice. Significance between three or more groups is determined by one way ANOVA test. Data are mean ± s.e.m.

**Extended Data Fig. 9.**
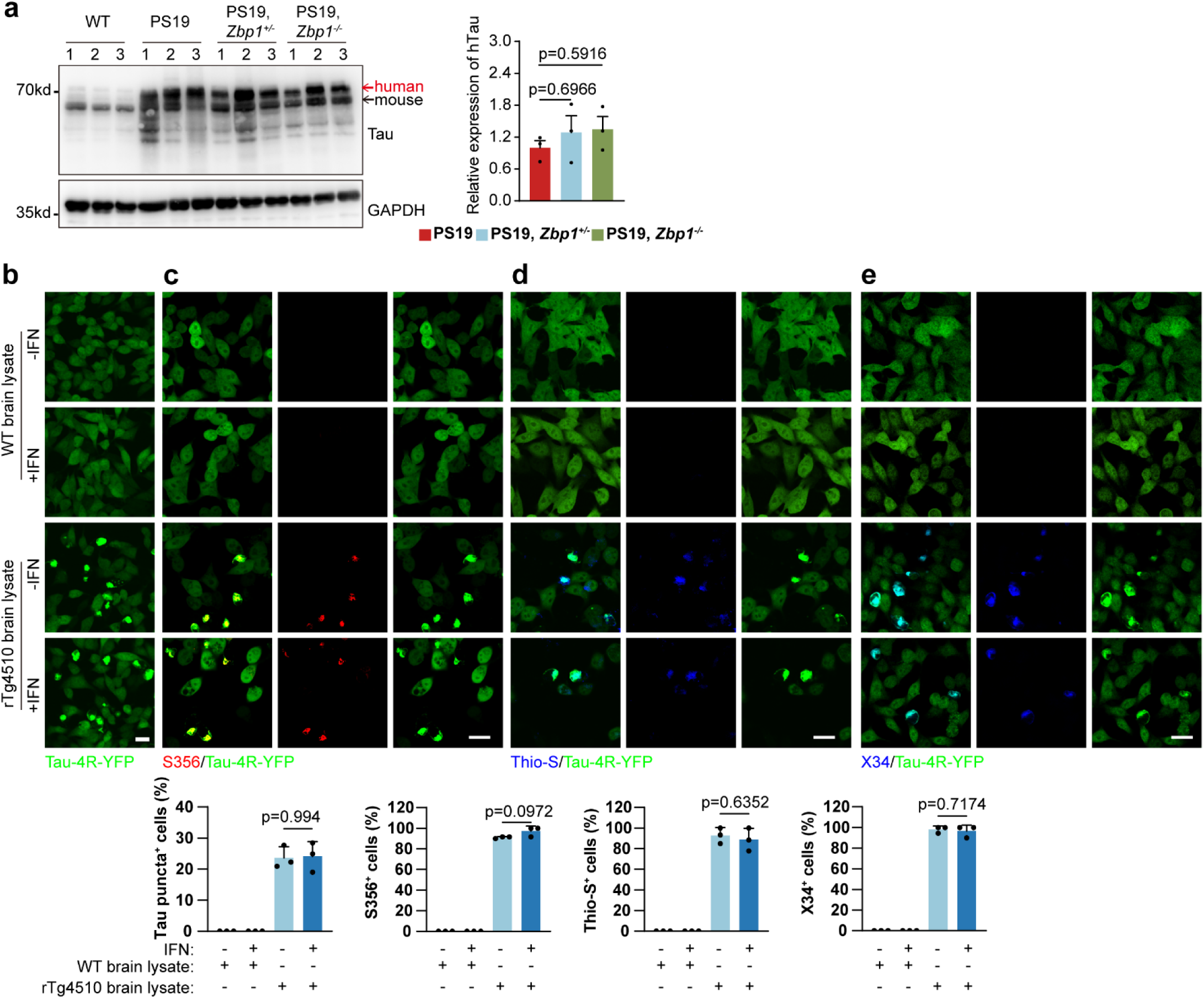
ZBP1 is not the direct upstream of Tau aggregates. **a.** Left: immunoblotting analysis of human Tau in the hippocampus from mice with indicated genotypes at 10-month-old. Right: quantification of human Tau protein level within indicated groups. n=3 mice. **b.** Upper: representative images of Tau puncta in SY5Y-Tau 4R cells treated with brain lysates from rTg4510 or WT mice with or without interferon stimulation (20 ng/ml IFNβ). Scale bar: 20 μm. Lower: quantification of Tau puncta within indicated groups. n=3 independent experiments. **c.** Upper: representative images of S356-positive cells treated with brain lysates from rTg4510 or WT mice with or without interferon stimulation (20 ng/ml IFNβ). Scale bar: 20 μm. Lower: quantification of S356-positive cells within indicated groups. n=3 independent experiments. **d.** Upper: representative images of Thio-S-positive cells treated with brain lysates from rTg4510 or WT mice with or without interferon stimulation (20 ng/ml IFNβ). Scale bar: 20 μm. Lower: quantification of Thio-S-positive cells within indicated groups. n=3 independent experiments. **e.** Upper: representative images of X34-positive cells treated with brain lysates from rTg4510 or WT mice with or without interferon stimulation (20 ng/ml IFNβ). Scale bar: 20 μm. Lower: quantification of X34-positive cells within indicated groups. n=3 independent experiments. Significance between three or more groups is determined by one way ANOVA test. Data are mean ± s.e.m.

